# *simplifyEnrichment*: an R/Bioconductor package for Clustering and Visualizing Functional Enrichment Results

**DOI:** 10.1101/2020.10.27.312116

**Authors:** Zuguang Gu, Daniel Hübschmann

## Abstract

Functional enrichment analysis or gene set enrichment analysis is a basic bioinformatics method that evaluates biological importance of a list of genes of interest. However, it may produce a long list of significant terms with highly redundant information that is difficult to summarize. Current tools to simplify enrichment results by clustering them into groups either still produce redundancy between clusters or do not retain consistent term similarities within clusters. We propose a new method named *binary cut* for clustering similarity matrices of functional terms. Through comprehensive benchmarks on both simulated and real-world datasets, we demonstrated that binary cut can efficiently cluster functional terms into groups where terms showed more consistent similarities within groups and were more mutually exclusive between groups. We compared binary cut clustering on the similarity matrices obtained from different similarity measures and found that the semantic similarity worked well with binary cut while similarity matrices based on gene overlap showed less consistent patterns. We implemented the binary cut algorithm in the R package *simplifyEnrichment* which additionally provides functionalities for visualizing, summarizing and comparing the clusterings.

## INTRODUCTION

Functional enrichment analysis or gene set enrichment analysis is a method widely used to evaluate whether genes of interest (GOI), *e*.*g*., differentially expressed genes, are over-represented or depleted in certain biological processes represented by gene sets. It is performed by testing the list of GOI against these gene sets *e*.*g*., through a hypergeometric test (1), by analyzing whether the gene sets are enriched at the top of a ranked gene list (2), or by evaluating the gene expression profile in individual gene sets through univariate or multivariate approaches (3). There are also methods to associate genomic regions of interest (ROI), *e*.*g*., regulatory domains, to gene sets, *e*.*g*., to identify which biological processes the ROIs might affect (4). Functional enrichment analysis depends on *a priori* biological knowledge encoded in pre-defined gene sets, which are obtained from multiple sources. Among them, Gene Ontology (GO) (5) is the most comprehensive source which organizes the biological terms in a hierarchical tree structure in the form of a Directed Acyclic Graph (DAG). Other popular sources are, for example, KEGG pathways (6) and MsigDB catalogues (7).

Enrichment results usually contain a long list of significantly enriched terms which have highly redundant information and are difficult to summarize, *e*.*g*., an analysis on 671 EBI Expression Atlas differential expression datasets (8) using GO gene sets with Biological Process ontology showed that there were 543 (80.9%) datasets having more than 250 significantly enriched GO terms under FDR < 0.05. The enrichment results can be simplified by clustering functional terms into groups where in the same group the terms provide similar information. The similarities between terms are important for the clustering. The calculation varies for different sources of gene sets. In the simplest and most general form, the similarity between two gene sets is based on gene overlap and is calculated as Jaccard coefficient, Dice coefficient or overlap coefficient (9). The Kappa coefficient is suggested to be more robust, and it is implemented in the widely used *DAVID* tool (10). For GO gene sets, there are more advanced methods which integrate the DAG structure to measure the semantic similarity between terms, *e*.*g*. the methods proposed in (11, 12) and implemented in the R package *GOSemSim* (13). The semantic similarity can also be calculated on other ontologies with DAG structure such as the Disease Ontology (14, 15).

One way to simplify the enrichment results is to cluster the similarity matrix of gene sets. Various clustering methods that have already been implemented in current tools can be applied, for example, hierarchical clustering (16), affinity propagation clustering (17) or density-based clustering (10). The similarity matrix can also be treated as an adjacency matrix and converted to a graph where the gene sets are the nodes and the similarity values are the weights of edges. Then, algorithms for detecting graph communities or modules can be applied for identifying the clusters. A popular tool is *Enrichment Map* (9).

Nevertheless, there are some common issues for clustering the similarity matrix of gene sets. Taking GO gene sets as an example, we call the root nodes *(i*.*e*., the three top terms Biological Process, Molecular Function and Cellular Component) *the root* of the GO tree. In the GO tree, there are branches with different depths, which causes two major problems for clustering. First, the size of GO clusters varies a lot, which means that there are large clusters which normally correspond to large branches inheriting from the root region of the GO tree, while at the same time, there are also many tiny clusters which correspond to the small branches in the very downstream of the GO tree. Due to the hierarchical relations in the tree, large clusters tend to have relatively low average similarities among terms while small clusters generally have relatively high average similarities. Methods such as *k*-means clustering cannot preserve the large clusters and instead split them into smaller clusters, which might still produce redundant information between clusters. Moreover, they are not able to separate tiny clusters while they prefer to merge them into one larger cluster. This over-segmentation and under-segmentation behaviour is mainly due to the fact that *k*-means clustering expects clusters to have similar sizes. Second, large GO clusters contain large amounts of terms and it is possible that a small fraction of terms in cluster A also share similarities to terms in cluster B, *i*.*e*., larger clusters are less mutually exclusive than smaller clusters, and this results in some graph community methods merging clusters A and B into one larger cluster due to the existence, albeit in small amounts, of terms shared by the two clusters. This in turn increases the inconsistency among gene sets in that merged cluster. Thus, an effective method is needed to balance between these two scenarios, reflecting generality and specificity of the GO system.

In this paper, we propose a new method named *binary cut* that clusters functional terms based on their similarity matrix. It recursively applies partition around medoids (PAM) with two groups on the similarity matrix and in each iteration step, a score is assigned to the submatrices to decide whether the corresponding terms should be further split into smaller clusters or not. We compared binary cut to a wide range of other clustering methods with 100 simulated GO lists and 468 GO enrichment results from real-world datasets, and we demonstrated that, with similarities calculated by semantic measurements, binary cut was more efficient to cluster GO terms such that inside clusters, terms shared more consistent similarity while at the same time the clusters had a higher degree of mutual exclusivity. Binary cut can also be applied to similarity matrices in general, *i*.*e*., based on gene overlap between gene sets and on any type of ontologies. However, the performance of binary cut varied, depending on the sources of gene sets. Similarity matrices based on gene overlap generally showed less consistent patterns than semantic similarity matrices and were not suggested to be used with binary cut.

We implemented binary cut in an R/Bioconductor package named *simplifyEnrichment*. After the functional terms have been clustered, *simplifyEnrichment* visualizes the summaries of clusters by word clouds, which helps users to easily interpret the common biological functions shared in the clusters. *simplifyEnrichment* can also export the clustering results to a web application so that users can interactively retrieve information of functional terms that belong to a certain cluster. Additionally, *simplifyEnrichment* provides a framework where user-defined clustering methods can be integrated and subsequently their results can be fed into the visualization functionality of *simplifyEnrichment*. Furthermore, *simplifyEnrichment* can compare multiple results from different clustering methods in a straightforward way.

## MATERIALS AND METHODS

### Similarity measure

If functional terms can be clustered into groups, the similarity matrices of corresponding gene sets are expected to represent a pattern where blocks can be observed on the diagonal where hierarchical clustering is applied on both rows and columns. Thus, efficiently clustering gene sets is equivalent to the task of identifying the *diagonal blocks* in the similarity matrix. The similarity between gene sets can be calculated as gene overlap by, *e*.*g*., the Jaccard coefficient, Dice coefficient or overlap coefficient. The three coefficients have similar forms. It has also been suggested to calculate the gene overlap by kappa coefficient which takes into account the possibility of gene occurrence in two gene sets just by chance (10). The formal definition of the four gene overlap coefficients can be found in [Supplementary file 1](https://simplifyenrichment.github.io/supplementary/suppl1_coefficient_definition/suppl1_coefficient_definition.html). If the ontologies have DAG structure such as GO, the topology information can be used to calculate the similarities, to generate a so-called *semantic similarity, e*.*g*., through information content-based approaches which measure the information each term obtains from its offspring terms. There exist a broad amount of algorithms for calculating semantic similarities, see (18) for a comprehensive overview.

As we observed from the benchmark datasets for GO terms, semantic similarities showed more distinct diagonal block patterns than similarity measures based on gene overlap (see comparisons in Section “General similarity by gene overlap”). The diagonal block pattern of the semantic matrix was frequently observed even for randomly sampled GO terms (see examples in [Supplementary file 2](https://simplifyenrichment.github.io/supplementary/suppl2_random_GO_examples/suppl2_random_GO_examples.html) and 4). Thus, we mainly use semantic similarity for clustering GO terms in this work. *simplifyEnrichment* uses the semantic measurements implemented in the R package *GOSemSim* (13) and takes the relevance method (12) as default (see comparisons of different semantic measurements in [Supplementary file 3](https://simplifyenrichment.github.io/supplementary/suppl3_examples_of_semantic_measures/suppl3_examples_of_semantic_measures.html)). Nevertheless, a similarity matrix by any other method and on any other ontology can be provided for an analysis with *simplifyEnrichment*.

### Clustering process

Let **M**_*a*_ and **M**_*b*_ denote the similarity matrices of two lists of GO terms, as illustrated in Figure 1. Intuitively, in this example GO terms in **M**_*a*_ have overall high pairwise similarities, thus they should be treated as one single cluster (Figure 1A), while terms in **M**_*b*_ show a two-block pattern on the diagonal and should be split into two sub-clusters (Figure 1B). A metric can be designed to decide whether a group of terms should be split or not. Based on this idea, we propose a new method named *binary cut* to cluster the similarity matrix of functional terms, executed in two phases:

**Figure 1.**
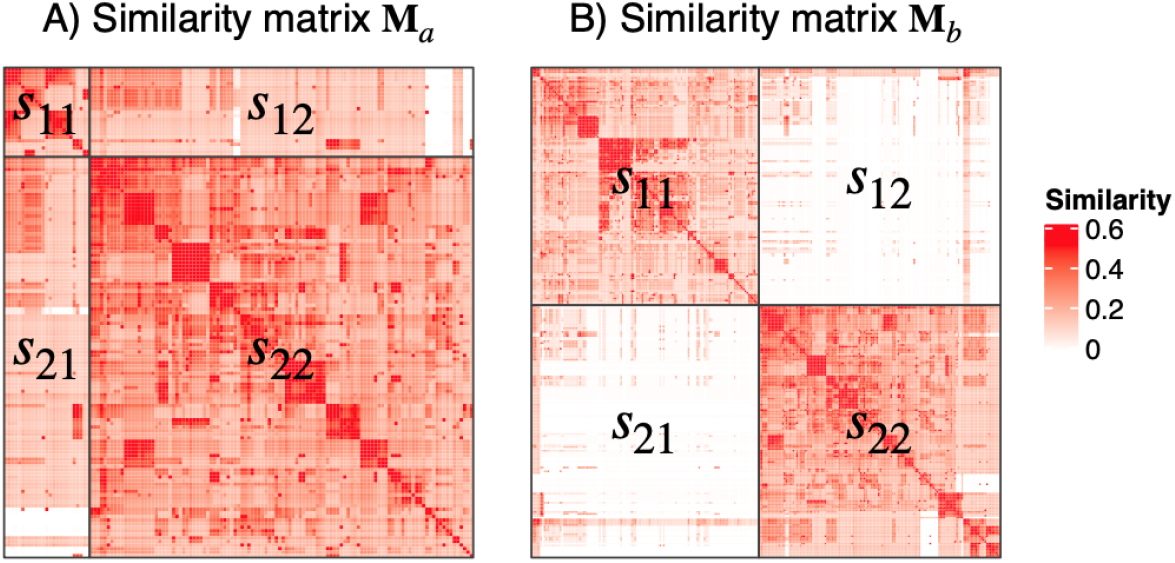
Examples of similarity matrices for two sets of GO terms, denoted as **M**_*a*_ and **M**_*b*_. Both matrices are split into two groups in the two dimensions. The values *s*_11_, *s*_12_, *s*_21_ and *s*_22_ are the medians of the entries in the corresponding submatrices. The values on the diagonal are excluded when calculating *s*_11_ and *s*_22_.

#### Phase 1: Applying divisive clustering and generating a dendrogram

This phase includes two steps:

##### Step 1

For the similarity matrix **M** of a given gene set, apply partitioning around medoids (PAM) with two-group classification on both rows and columns, which partitions **M** into four submatrices, denoted as **M**_11_, **M**_12_, **M**_21_, and **M**_22_ where the indices represent the groups in the two matrix dimensions. Next calculate the following scores *s*_11_, *s*_12_, *s*_21_ and *s*_22_ where *s*_11_ and *s*_22_ are the median values of all entries in **M**_11_ and **M**_22_ excluding the diagonals. *s*_12_ and *s*_21_ are the median values of all entries in **M**_12_ and **M**_21_. Since the similarity matrix is always symmetric, *s*_12_ and *s*_21_ are equal. *s*_11_ or *s*_22_ is defined to be 1 if **M**_11_ or **M**_22_ have only one row. We then define the score *s* as

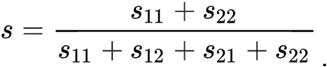

The diagonal blocks **M**_11_ and **M**_22_ correspond to similarity measures within sub-clusters, whereas the off-diagonal blocks **M**_12_ and **M**_21_ correspond to similarity measures between sub-clusters. In a successful clustering, similarity measures will be higher within clusters than between clusters, and this roughly translates to the median values *s*_11_, *s*_12_, *s*_21_ and *s*_22_, which results in *s*_11_+ *s*_22_ ≥ *s*_12_+ *s*_21_. Thus, the values of *s* approximately range between 0.5 and 1. If the similarity matrix **M** shows overall similarities, *s* is close to 0.5 (Figure 1A) and it should not be split any more, while if the GO terms can still be split into more groups, *s*_12_ and *s*_21_ are close to 0, which results in *s* close to 1 (Figure 1B).

##### Step 2

Apply Step 1 to the two submatrices **M**_11_ and **M**_22_ respectively. The clusterings in Steps 1 and 2 are executed recursively and the process is saved as a dendrogram where the score *s* is attached to every node in the dendrogram which corresponds to every submatrix in the iteration. The clustering stops when the number of terms in a group reaches 1; these are the leaves of the dendrogram.

#### Phase 2: Cutting the dendrogram and generating clusters

Since every node in the dendrogram has a score *s* computed in Phase 1, *s* is simply compared to a cutoff (0.85 as the default; it can be optimally selected by *simplifyEnrichment*). If *s* is larger than the cutoff, the two branches from the node are split, else all the terms under the node are taken as a single cluster.

Nodes with large numbers of terms tend to have relatively smaller *s*, thus, it is possible that at a certain node, *s* does not exceed the cutoff but is very close to it, while its child nodes have values of *s* larger than the cutoff. In this case, we don’t want to close the node so early and we still split this node into two subgroups so that its child nodes can be split furthermore. Thus, the rule in Phase 2 is modified as follows: if the score *s* of a given node does not exceed the cutoff but it is larger than 0.8*cutoff, the node is still split if at least one of its child nodes has a score which exceeds the cutoff. This is equivalent to reassigning the maximal scores of its two child nodes to the score *s* of this given node.

In the iterations of binary cut clustering, there might exist more than two diagonal blocks in the submatrices. Iterative application of bifurcations, *i*.*e*., two-group partitioning, can resolve any number of underlying diagonal blocks or clusters. As we have broadly tested on a large number of datasets, in most cases, PAM kept the completeness of diagonal blocks based on the two-group classification.

An example of the process of binary cut clustering is demonstrated in Figure 2. Figure 2A-C illustrate the clusterings in the first three iterations. Figure 2D illustrates the final dendrogram where the nodes that are split are marked with crosses. To optimize the clustering process, *simplifyEnrichment* supports partial clustering to reduce the computing time where the complete dendrogram is not generated and the bifurcation stops on a node as long as the corresponding *s* does not exceed the cutoff.

**Figure 2.**
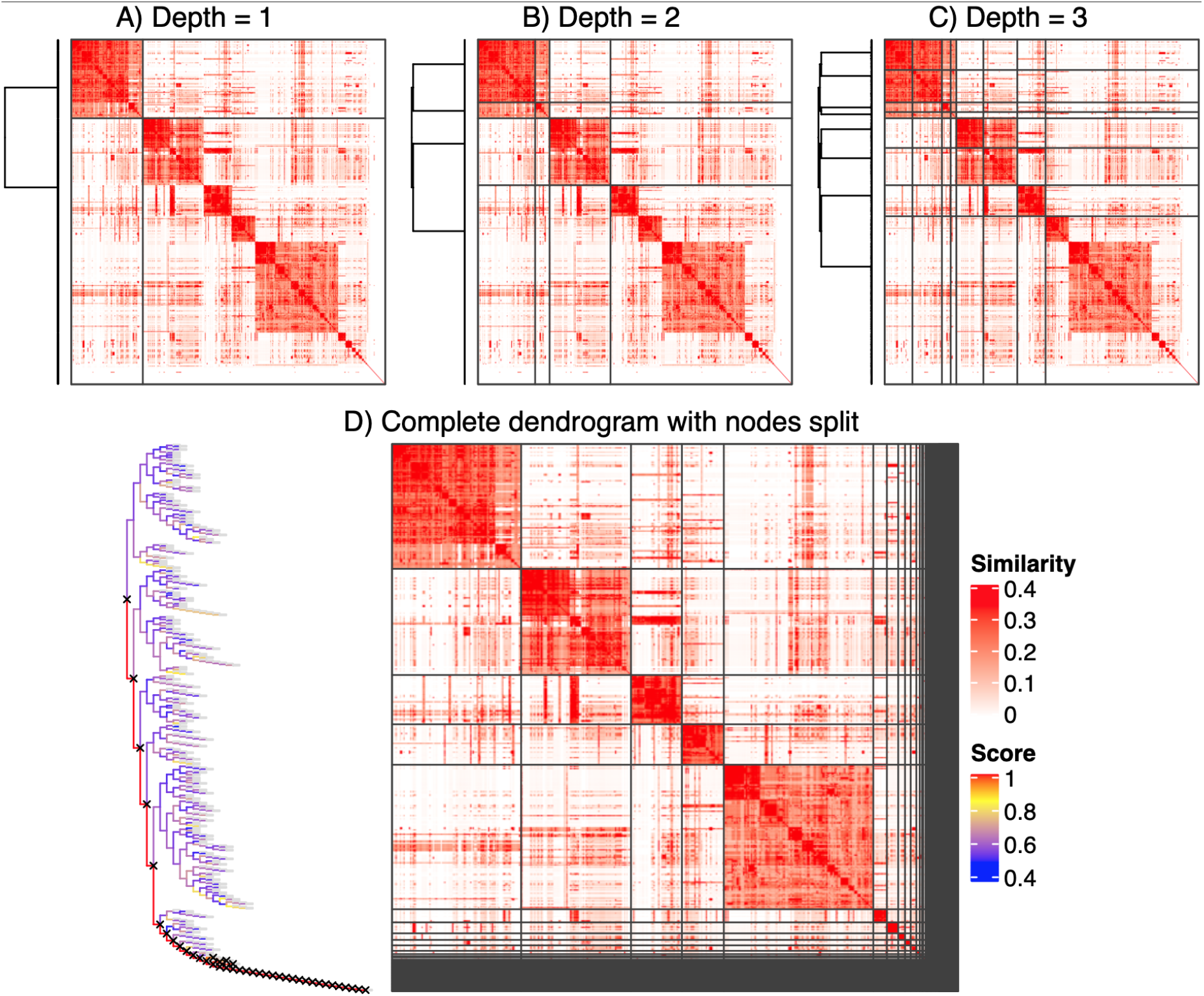
A demonstration of the binary cut clustering with 500 random GO terms. A-C) The clusterings in the first three iterations; D) The complete dendrogram from binary cut clustering. The colors of the dendrogram segments correspond to the scores *s* assigned to the nodes. Nodes to split are marked with crosses.

### Software implementation

The binary cut algorithm is implemented in an R/Bioconductor package named *simplifyEnrichment*. The input of the package is a similarity matrix that is provided either directly by the user or by the functions implemented in *simplifyEnrichment*, such as the function `GO_similarity()` for calculating GO semantic similarity or the function `term_similarity()` that measures general gene set similarity by Jaccard coefficient, Dice coefficient, overlap coefficient or kappa coefficient based on gene overlap.

The function `simplifyGO()` performs clustering for GO terms and visualizes the results. Once the GO terms have been clustered, the biological descriptions for the terms are automatically extracted. The summaries of the biological functions in clusters are visualized as word clouds and are attached to the similarity heatmap, which gives a direct illustration of the common biological functions involved in each cluster (Figure 3). The function `simplifyEnrichment()` performs clustering on similarity matrices from any type of ontology. The static heatmap demonstrated in Figure 3 can be exported to a Shiny web application by the function `export_to_shiny_app()` so that users can interactively select GO terms that belong to a specific cluster from the heatmap for deeper exploration (see example in [Supplementary file 11](https://simplifyenrichment.github.io/supplementary/suppl11_shiny_app/suppl11_shiny_app.html)).

**Figure 3.**
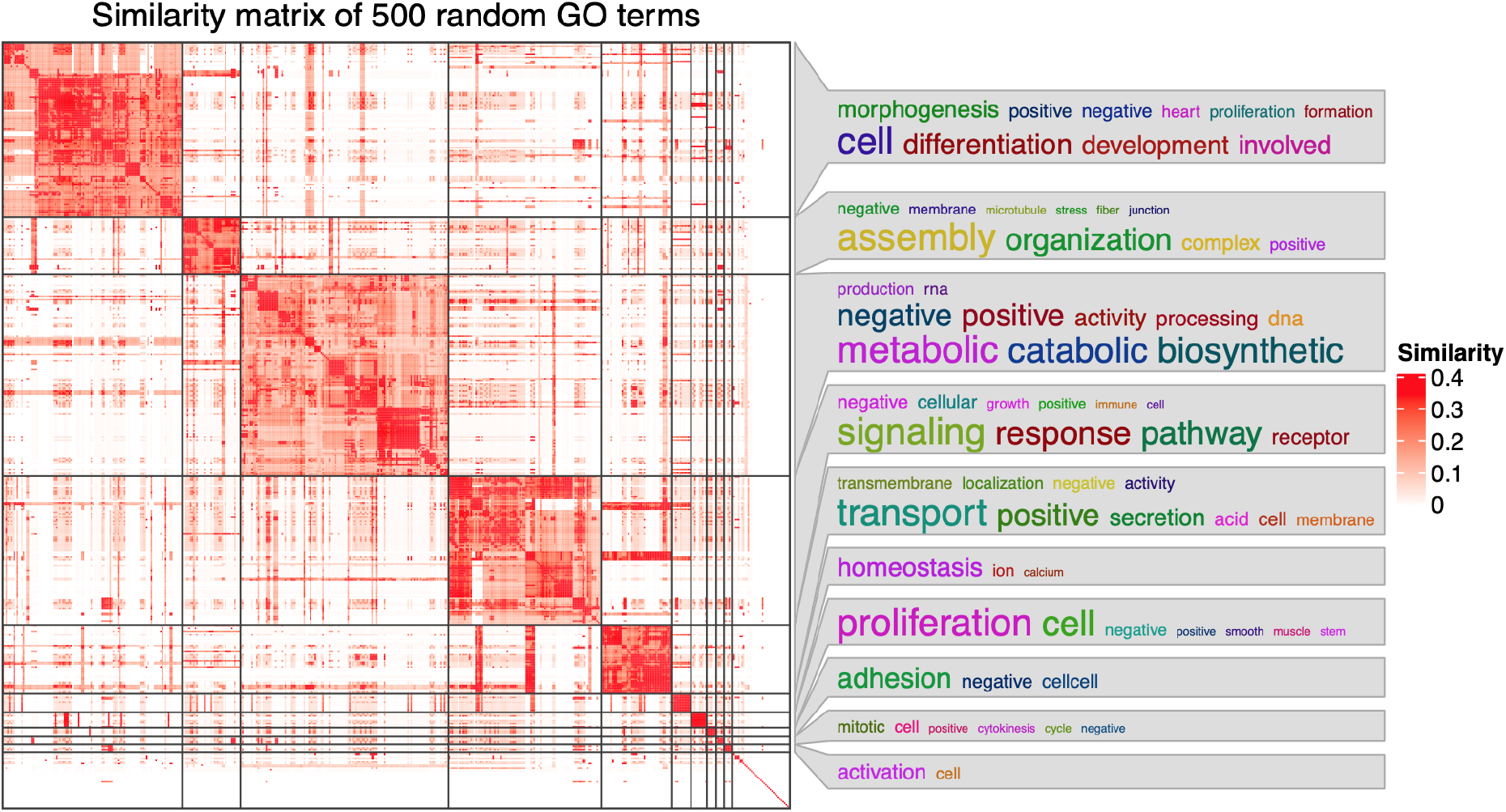
Example of a similarity heatmap from 500 random GO terms that have been clustered and annotated with word clouds. The bottom right cluster with no word cloud annotation contains all other small clusters with numbers of terms less than 5. The plot was made by the function *simplifyGO()*.

*simplifyEnrichment* also allows to integrate other clustering methods. New clustering functions can be added by the function `register_clustering_methods()`. The function `compare_clustering_methods()` applies multiple clustering methods on the same similarity matrix and compares them via heatmaps (see example in Figure 4).

**Figure 4.**
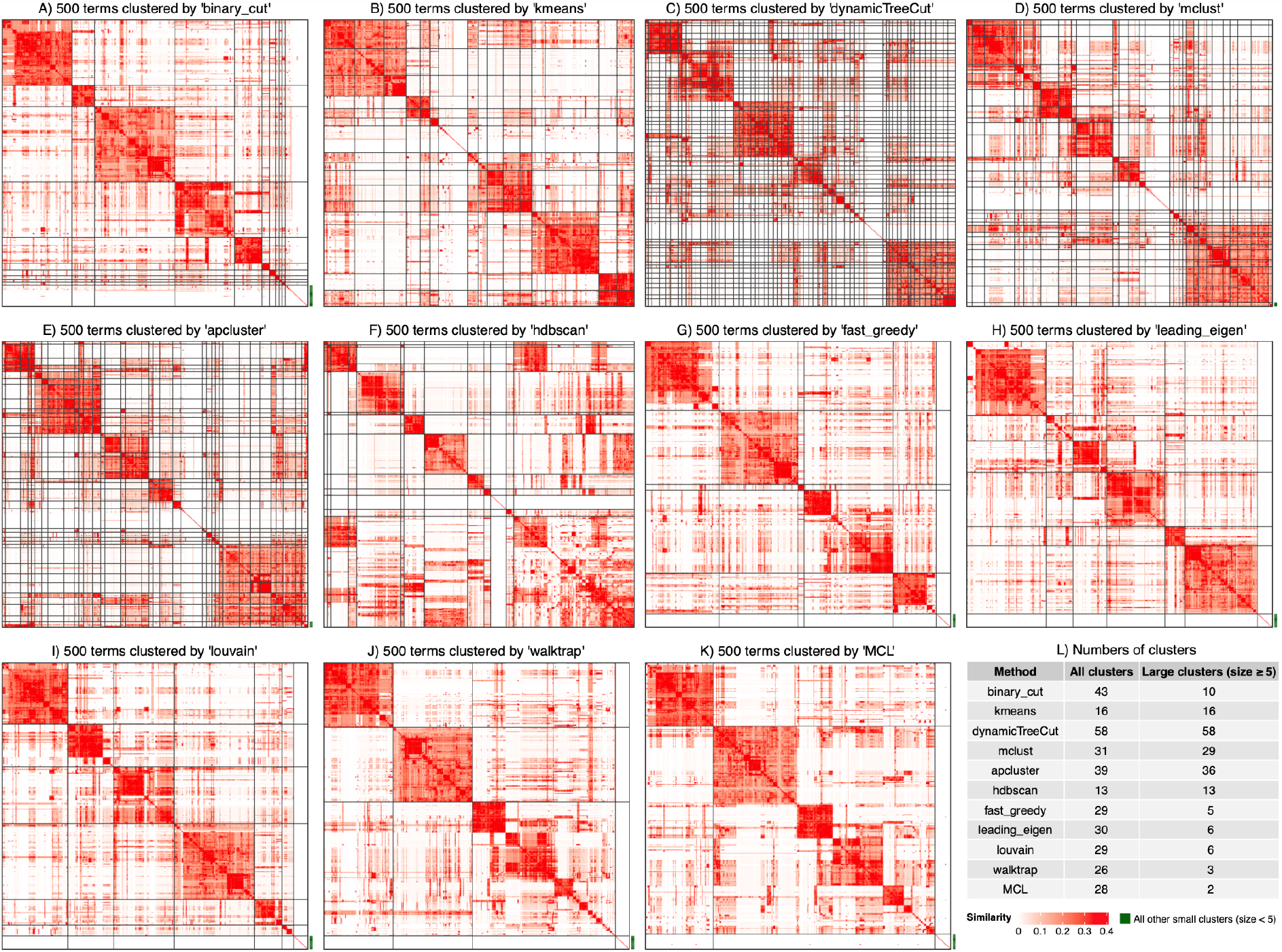
Comparison of different clustering methods. A-K) Clusterings from 11 different methods. L) Numbers of all clusters and numbers of large clusters with size ≥ 5. For some methods, the small clusters (size < 5) were put into one single cluster on the bottom right of the heatmap and were marked by green lines. All the methods were applied to the same GO semantic similarity matrix from 500 random GO terms from the Biological Process ontology. The plots were generated by the function *compare_clustering_methods()*.

## RESULTS

We compared the following 10 clustering methods to binary cut: *k*-means clustering (kmeans) where the optimized number of clusters was selected based on the distribution of within-cluster sum of squares (WSS), dynamicTreeCut (19), mclust (20), apcluster (21), hdbscan (22), the following graph community methods implemented in the R package *igraph* (23): *fast greedy, leading eigen, louvain and walktrap, as well as one additional community method MCL (24) which was recently used in an implementation for clustering GO terms (25). These selected methods are based on various categories of clustering algorithms and are by default supported in simplifyEnrichment*. GO semantic similarities were calculated by the relevance method (12) with the R package *GOSemSim*.

### Application of *simplifyEnrichment* to random GO lists

500 random GO terms were uniformly sampled from the Biological Process (BP) ontology. Clusterings of 11 different methods are illustrated in Figure 4. dynamicTreeCut, mclust and apcluster generated huge numbers of small clusters (Figure 4C-E). kmeans and hdbscan generated intermediate numbers of clusters, but kmeans failed to preserve large clusters even if overall similarities within them were high (*e*.*g*., the first three clusters in Figure 4B), while for hdbscan, the largest cluster did not show consistent similarities for all term pairs and it should be split further (the bottom right cluster in Figure 4F). Community methods of fast greedy, leading eigen, louvain, walktrap and MCL identified both small and large clusters; however, some large clusters may still be split further (Figure 4G-K), as most strikingly visualized for MCL where only two large clusters were identified (Figure 4K). In comparison, binary cut generated clean clusters and it was able to identify large and small clusters at the same time (Figure 4A).

We then benchmarked the clustering methods quantitatively. To this end, the random GO lists of 500 BP terms were generated 100 times and the following metrics were used:

#### Difference score

It measures the difference of the similarity values of the terms within clusters and between clusters. For a similarity matrix denoted as **M**, and for term *i* and *j* where *i* ≠ *j*, the similarity value *x*_*i, j*_ is saved to the vector **x**_1_ when term *i* and *j* are in the same cluster. *x*_*i, j*_ is saved to the vector **x**_2_ when term *i* and *j* are in different clusters. The difference score measures the distribution difference between **x**_1_ and **x**_2_, calculated as the Kolmogorov-Smirnov statistic between the two distributions of **x**_1_ and **x**_2_.

#### Number of clusters

For each clustering, there are two numbers: the total number of clusters and the number of clusters with size ≥ 5 (*i*.*e*., the large clusters). The two values can be used to test whether the clustering methods can identify small clusters.

#### Block mean

Mean similarity values of the diagonal blocks in the similarity heatmap. Using the same convention as for the difference score, the block mean is the mean value of **x**_1_.

As shown in Figure 5A, binary cut had the highest difference score, reflecting that clusterings obtained with binary cut had the most distinct differences in similarity values within clusters and between clusters, therefore the clusters identified by binary cut were the most mutually exclusive among all methods. dynamicTreeCut and apcluster generated a huge number of clusters (Figure 5B, examples in Figure 4C and 4E). These two methods can be considered too stringent for clustering GO terms, in addition there still are high levels of redundant information among different clusters. In comparison, graph community methods and binary cut generated moderate numbers of clusters, and especially the numbers of “large clusters” (size ≥ 5) dropped dramatically compared to the total numbers of clusters, indicating that these methods were able to identify small clusters. Binary cut generated clusters with moderate block mean values (Figure 5C, examples in Figure 4G-H). In comparison, kmeans, dynamicTreeCut, mclust and apcluster generated high block mean values, reflecting that they were not able to preserve large clusters with intermediate similarities. Graph community methods generated low block mean values, mainly due to the fact that terms in the clusters were not necessary to have high similarities to all other terms (examples in Figure 4G-H). The method hdbscan generated intermediate numbers of clusters and had intermediate block mean values, but it had the lowest difference score in the comparisons, which implies that differences between similarities within clusters and between clusters were small (example in Figure 4F). In conclusion, these comparisons indicated that binary cut kept both generality and specificity of GO clusters. The analysis reports for all 100 random GO lists can be found in [Supplementary file 4](https://simplifyenrichment.github.io/supplementary/suppl4_examples/suppl4_examples.html).

**Figure 5.**
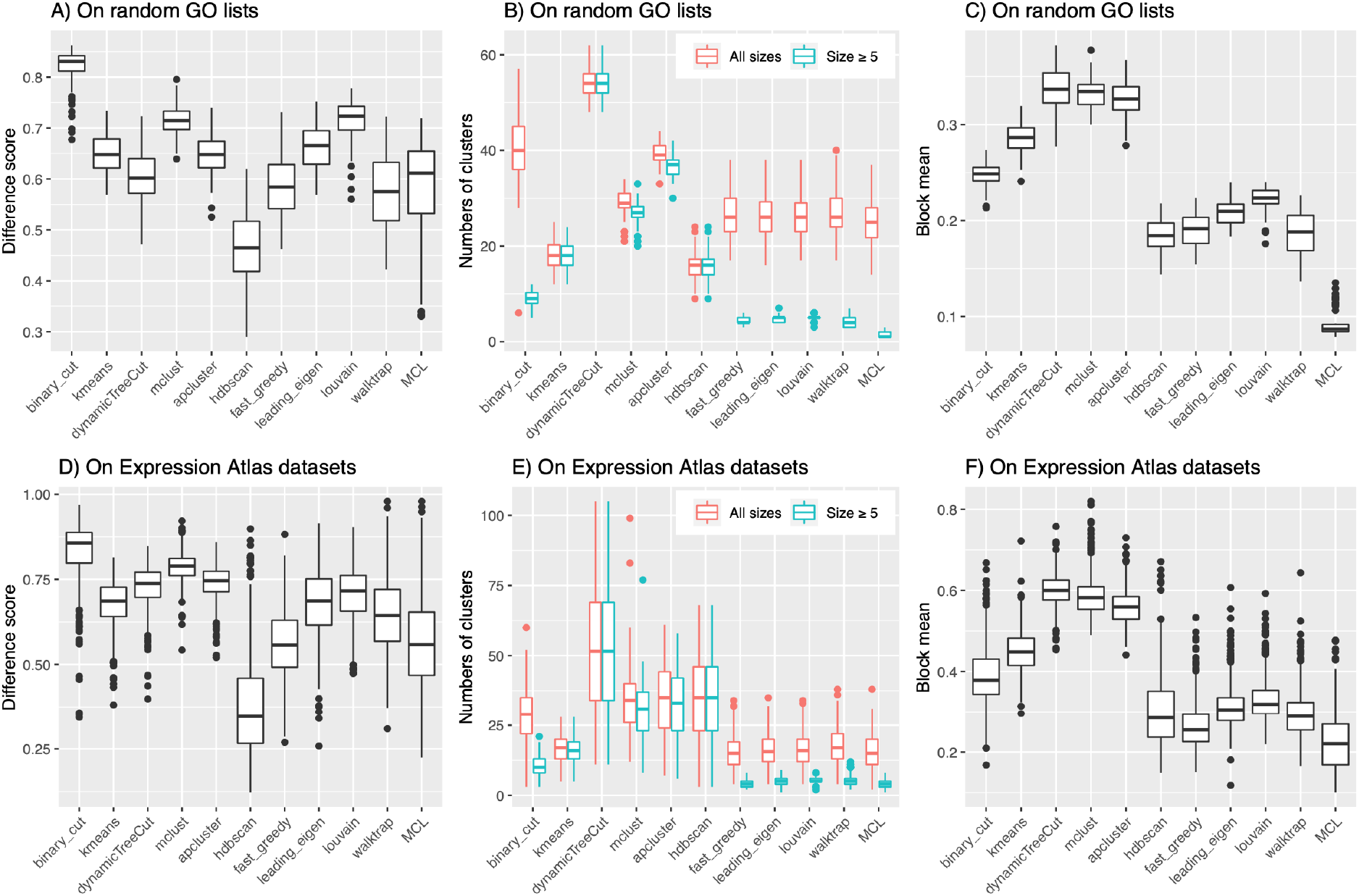
Benchmarks of different clustering methods. A-C) Difference scores, numbers of clusters and the block mean values for the clusterings identified by 11 methods. The analysis was applied to 100 random GO lists of 500 BP terms. D-F) Similar as Figure A-C, but on the functional enrichment results from 468 Expression Atlas datasets.

Of note, the similarity matrices of GO terms based on semantic measures showed clear diagonal block patterns even for lists of random GO terms. In [Supplementary file 3](https://simplifyenrichment.github.io/supplementary/suppl3_examples_of_semantic_measures/suppl3_examples_of_semantic_measures.html), we compared the five semantic similarity measurements supported in the *GOSemSim* package and found that the methods “Rel” (*i*.*e*., the relevance method) (12), “Resnik” (26) and “Lin” (27) produced similar and clear diagonal block patterns. They are thus suitable for use with binary cut clustering. Furthermore, an analysis of the complete set of terms of GO BP ontology showed that with semantic similarity, globally the similarity matrix had diagonal block patterns where the clusters corresponded to several top functional categories ([Supplementary file 7](https://simplifyenrichment.github.io/supplementary/suppl7_global_GO_similarity/suppl7_global_GO_similarity.html)). Uniformly sampling from all BP terms tends to retain these modular patterns.

By default and in every iteration step, *simplifyEnrichment* partitions the current submatrix into two groups using PAM. In [Supplementary file 8](https://simplifyenrichment.github.io/supplementary/suppl8_partitioning_methods/suppl8_partitioning_methods.html), we compared binary cut clustering on 100 random GO BP lists with the following three partitioning methods: PAM, *k*-means++ and hierarchical clustering with the “ward.D2” method. *k*-means++ is an optimized *k*-means clustering by pre-selecting proper initial centroids (28, 29). The results showed that the three partitioning methods performed very similarly, *i*.*e*., with similar difference scores, numbers of clusters and block mean values among 100 random GO lists. The final clusterings of GO terms highly agreed (on average 89% agreement, [Supplementary file 8](https://simplifyenrichment.github.io/supplementary/suppl8_partitioning_methods/suppl8_partitioning_methods.html): Figure S8.1) between the three partitioning methods. The three methods are all supported in *simplifyEnrichment* and they can be selected specifically for individual datasets. *simplifyEnrichment* also supports running all three methods simultaneously for an analysis and automatically selecting the method that generates clustering with the highest difference score.

### Application of simplifyEnrichment to real-world datasets

To assess the performance of *simplifyEnrichment* also on actual biological data, we analyzed all datasets from the Expression Atlas (8) that have differential expression analysis results. We applied functional enrichment analysis with the R package *clusterProfiler* (30) to the significantly expressed genes (FDR < 0.05) with the GO Biological Process ontology. We only took those datasets for which the number of significant genes was in the interval [500, 3000] and the number of significant GO terms (FDR < 0.05) was in [100, 1000]. This yielded 468 GO lists. Figure 5D-F illustrate the results for the comparisons among the 11 clustering methods. The conclusion was very similar to the benchmark with random GO datasets: binary cut outperformed other clustering methods. The analysis reports for 468 individual Expression Atlas datasets can be found in [Supplementary file 4](https://simplifyenrichment.github.io/supplementary/suppl4_examples/suppl4_examples.html).

We mainly benchmarked GO term enrichment with the BP ontology, which is the major ontology category in GO (65.9% of all GO terms). In [Supplementary file 4](https://simplifyenrichment.github.io/supplementary/suppl4_examples/suppl4_examples.html), we also applied binary cut to random GO lists from Cellular Component (CC) and Molecular Function (MF) ontologies. The results were very similar to those obtained with BP terms where binary cut outperformed other clustering methods on the semantic similarity matrices.

### General similarity by gene overlap

Many ontologies (*e*.*g*. MsigDB gene sets) are only represented as lists of genes for which the similarities are merely measured by gene overlap. While in the previous section, the benchmark was based on only one similarity measure, *i*.*e*., semantic similarity, we thus performed a second benchmark and compared the performance of binary cut when applied to similarity matrices calculated by different similarity measures. In addition to semantic similarity, these included four gene overlap measures: Jaccard coefficient, Dice coefficient, overlap coefficient and kappa coefficient. For this second benchmark, we again used the 100 random GO lists with 500 BP terms as well as 468 significant GO lists from Expression Atlas datasets.

When GO terms were randomly generated, with similarities measured by Jaccard coefficient, Dice coefficient and kappa coefficient, 500 terms were split into large numbers of clusters (on average 358 clusters, Figure 6A, red boxes), where on average, less than two terms were in a single cluster (Figure 6B, red boxes). The numbers of clusters dropped dramatically if only counting clusters with size ≥ 5 (on average 6 clusters for all five similarity measures, Figure 6A, blue boxes). For the clusters with size ≥ 5, the average numbers of GO terms per cluster were 20 with the Jaccard coefficient, 17 with the Dice coefficient and 12 with the kappa coefficient, while 53 were identified with semantic measurement (Figure 6B, blue boxes). For the clusters with size < 5 which we defined as “small clusters”, the Jaccard coefficient generated on average 375 small clusters (covering 84.5% of all terms), the Dice coefficient generated 335 small clusters (81.4% of all terms) and the kappa coefficient generated 345 small clusters (84.9% of all terms), while in comparison, semantic measurement only generated 31 small clusters (7.8% of all terms) (see difference between red and blue boxes in Figure 6A). This implies that binary cut generated huge number of small clusters when based on the similarity matrices of Jaccard coefficient, Dice coefficient and kappa coefficient and that these three coefficients could not assign similarities for most of the term pairs, thus they were not efficient for clustering. Examples of the clusterings under different similarity measures can be found in Figure 6D-E and [Supplementary file 5](https://simplifyenrichment.github.io/supplementary/suppl5_compare_similarity_random_GO/suppl5_compare_similarity_random_GO.html). The overlap coefficient performed differently from the other three gene overlap measures (32.3% agreement of the clusterings, Figure 6C). In 30% of all 100 random GO lists, the 500 GO terms could not even be separated and the numbers of clusters for them were only identified as one ([Supplementary file 5](https://simplifyenrichment.github.io/supplementary/suppl5_compare_similarity_random_GO/suppl5_compare_similarity_random_GO.html): Figure S5.2-S5.4). The individual heatmaps of overlap similarities showed that in most cases only one major cluster was generated with a marginal pattern in which a small amount of terms showed high similarities to most of the other terms and no diagonal block pattern was observable ([Supplementary file 5](https://simplifyenrichment.github.io/supplementary/suppl5_compare_similarity_random_GO/suppl5_compare_similarity_random_GO.html): Table S5.1), thus binary cut had very weak performance on it. The marginal pattern is due to the definition of the overlap coefficient according to which the score was normalized by the size of the smaller gene set. In such a setting, a term located in the downstream part of the GO tree, *i*.*e*., close to the leaves, has the same overlap coefficients as all its ancestor terms because parent terms include all genes of their child terms. The heatmaps for individual datasets ([Supplementary file 5](https://simplifyenrichment.github.io/supplementary/suppl5_compare_similarity_random_GO/suppl5_compare_similarity_random_GO.html): Table S5.1) showed that, for the randomly sampled GO lists, similarity values calculated by gene overlap were very weak and noisy, which led to the fact that in most cases, terms could not be clustered. In comparison, the semantic similarity matrices generated intermediate numbers of clusters and had clear diagonal block patterns (Figure 6D).

**Figure 6.**
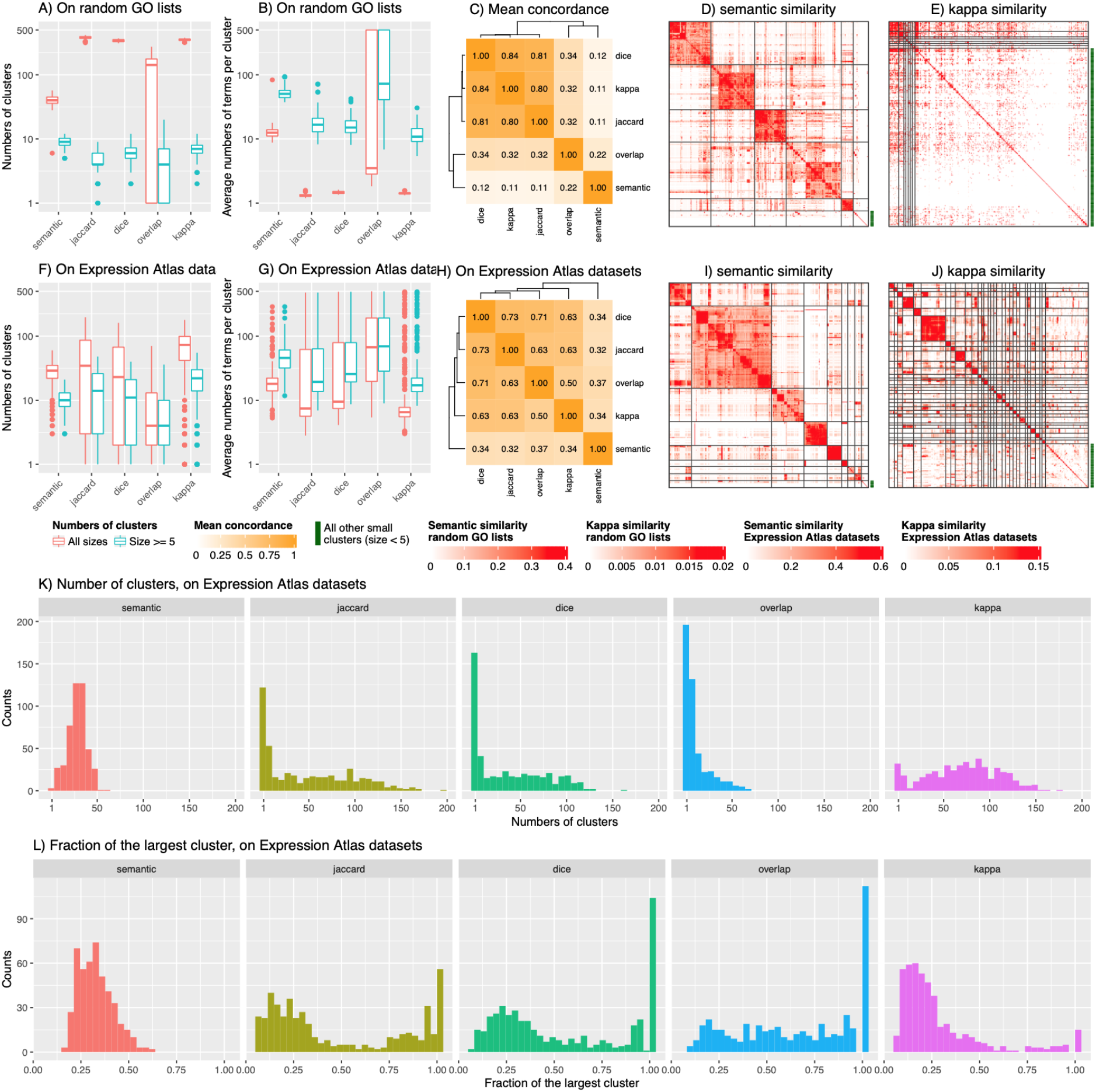
Comparison of clusterings on similarity matrices by different similarity measures. A) Numbers of clusters. B) Average numbers of terms per cluster. Y-axes in A and B are in log10 scale. C) Mean concordance of the clusterings. The definition of concordance can be found in [Supplementary file 10](https://simplifyenrichment.github.io/supplementary/suppl10_concordance/suppl10_concordance.html). The analysis was applied to 100 random GO lists with 500 BP terms. D-E) Examples of binary cut clustering on two similarity matrices from the list of random GO terms by semantic and kappa measurements. F-J) Analogous to A-E, but on the functional enrichment results from 468 Expression Atlas datasets. I and J are based on the same dataset. K) Distributions of cluster sizes on similarity matrices from different similarity measures. L) Distributions of the fraction of the largest cluster. K and L are based on Expression Atlas datasets.

Analysis on the Expression Atlas datasets with GO BP ontology also showed that binary cut applied to similarity matrices obtained by semantic similarity generated very different clusterings from the four gene overlap similarity matrices (Figure 6F-J). The cluster sizes obtained from the matrices measured by Jaccard coefficient, Dice coefficient and kappa coefficient had very broad ranges where many datasets had more than 100 clusters (on average 16.3% of all datasets, Figure 6K, an example in Figure 6J). On the other hand, with Jaccard coefficient, Dice coefficient and overlap coefficient, binary cut clustered terms only into small numbers of clusters for many datasets (less than 5 clusters in on average 39.7% of all datasets, Figure 6K). In comparison, semantic similarity generated numbers of clusters in a moderate range (14 to 40 for the 10th to the 90th percentile, Figure 6K). Moreover, this analysis showed that the largest cluster comprised more than 80% of all terms in 30.6% of the datasets when using the Jaccard coefficient, in 33.8% of the datasets when using the Dice coefficient and in 39.7% of the datasets when using the overlap coefficient (Figure 6L). This implies that these three coefficients cannot efficiently separate terms in many datasets. Taken together, semantic similarity matrices worked better than gene overlap similarities on the real-world datasets.

Besides GO gene sets, we also applied other ontologies to Expression Atlas datastes, *i*.*e*., gene sets from DO, KEGG, Reactome and MsigDB, only with gene overlap as similarity measures ([Supplementary file 4](https://simplifyenrichment.github.io/supplementary/suppl4_examples/suppl4_examples.html)). We found that only the similarity matrices based on gene overlap coefficients of Reactome and MsigDB C4 gene sets can be used for binary cut clustering while the similarity matrices from other gene sets generally showed low consistency between terms and did not have clear diagonal block patterns, thus, they are not suited for application of binary cut.

### Comparison of enrichment results from multiple lists of genes - a case study

It is a common task to integrate functional enrichment results from multiple lists of genes into one plot, *e*.*g*., the enrichment results from up-regulated and down-regulated genes in a two-group differential expression analysis. One of the frequently used methods is to make barplot-like graphics where heights of bars correspond to -log10(p-value) or -log10(FDR) and the enrichment of the up-regulated and down-regulated genes are assigned with positive and negative signs respectively. This method is limited because users need to pre-select functional terms only to a small number either by selecting top n terms with the highest significance or by manually picking representative terms that have distinct biological meanings. The former way might lose important terms with less significance and the latter might be easily affected by subjectivity. Also this is limited only to two gene lists. Here, we used a public dataset to demonstrate a strategy to efficiently compare enrichment results from multiple gene lists without losing any information, which makes it easy to detect, *e*.*g*., the biological functions that are uniquely enriched only in one gene list. It works for arbitrary numbers of gene lists. This strategy was implemented in the function `simplifyGOFromMultipleLists()` in *simplifyEnrichment*.

Figure 7A illustrates a heatmap of signature genes from a three-group classification of the Golub leukemia dataset (31, 32). The classification was obtained by consensus partitioning with the R package *cola* (33). *The signature genes were additionally clustered into three groups by k*-means clustering and the groups were labelled as “km1”, “km2” and “km3”. GO enrichment analysis was applied to the three groups of genes separately with the R package *clusterProfiler* (30). To compare the enrichment results from the three gene lists, binary cut was applied directly to the union of the three significant GO term lists (FDR < 0.01) and a heatmap of FDR was put on the left of the GO similarity heatmap to visualize whether the GO terms were significant in the corresponding gene list (Figure 7B). This strategy keeps all significant GO terms without removing anyone. Also it allows more specific studies of how the enrichment varies between gene lists. For example, in Figure 7B, it can be observed that genes down-regulated in acute myeloid leukemia (AML) samples (km1 group) were more specifically enriched in cell cycle and metabolic processes (labelled with “1” in Figure 7B), that genes up-regulated in AML (km2 group) were more specifically enriched in lymphocyte activation (labelled with “2” in Figure 7B), and that genes down-regulated in a subset of acute lymphoblastic leukaemia (ALL) samples (km3 group) were more specifically enriched in cell differentiation (labelled with “3” in Figure 7B).

**Figure 7.**
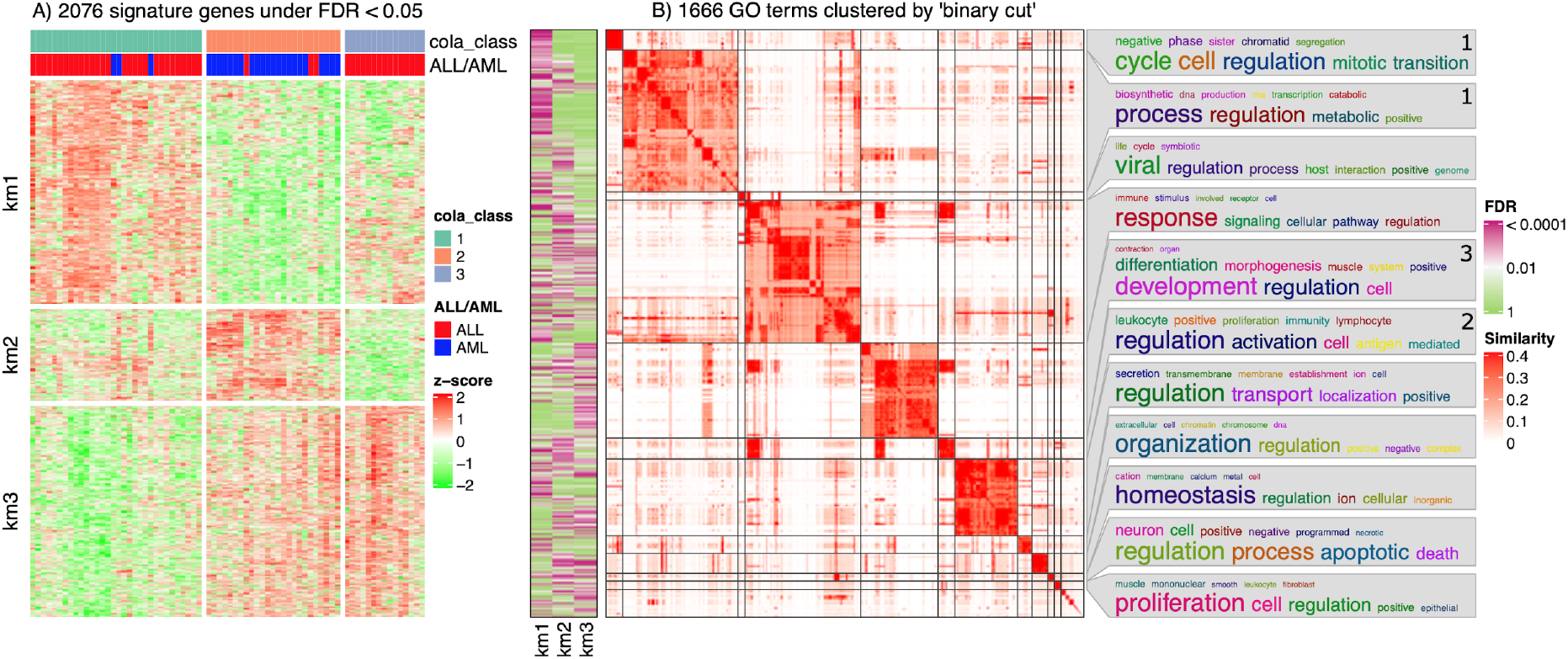
Compare enrichment results from three gene lists. A) Heatmap of the expression of signature genes from a three-group classification of Golub leukemia dataset. The three gene lists were generated by applying *k*-means clustering on rows of the expression matrix. B) GO terms that were significant in any enrichment results of the three gene lists were clustered and their similarities were visualized as a heatmap. The left heatmap demonstrates whether the GO terms were significant in the corresponding gene list. The labels of “1, 2 and 3” on word clouds are explained in the main text.

This strategy had also been applied in a previous study (33) where it compared biological functions of gene lists generated by four different feature selection methods for consensus partitioning to assess which feature selection method generates more biologically meaningful features.

## DISCUSSION

Simplifying functional enrichment results is necessary for easy interpretation without missing important biological terms. The essential step to simplify enrichment results is to cluster the functional terms; later downstream analysis can be applied to summarize the major biological functions in each cluster, such as extracting the most significant terms in each cluster (30) or looking for a minimal subset of terms that cover genes from all terms in that cluster (34). A proper clustering method should identify groups of terms that provide more distinct separation between clusters and reduce large numbers of terms into an amount easier to read and analyze. In this work, we developed the R package *simplifyEnrichment* which uses a new method called *binary cut* to efficiently cluster functional terms and to summarize the general functions in the obtained clusters by word cloud visualizations.

Binary cut is based on the observation that clustering gene sets efficiently is equivalent to the task of identifying *diagonal blocks* in the similarity matrix of these gene sets after hierarchical clustering of both rows and columns. The algorithm is implemented in two phases: (1) applying divisive clustering and generating a dendrogram on both rows and columns, and (2) cutting the dendrogram and generating clusters. By default, phase (1) relies on recursive application of partitioning around medoids (PAM) with two-group classification, but other partitioning methods, including *k*-means++ and hierarchical clustering with the “ward.D2” method, also performed well.

In the binary cut clustering process, a score is calculated based on the two-group partitioning on the current submatrix and it is compared against a cutoff to decide whether the corresponding terms are treated as a single cluster or should be further split. In most cases, the default cutoff is robust and works well if a diagonal pattern is observable in the similarity matrix. Nevertheless, there are still cases where manual adjustment on the cutoff needs to be applied, such as a large cluster which is composed of several small and dense subclusters. A typical example is the clustering on the semantic similarity matrices of Disease Ontology terms ([Supplementary file 4](https://simplifyenrichment.github.io/supplementary/suppl4_examples/suppl4_examples.html)). The cutoff can be fine-tuned according to what levels of similarities users want to keep. *simplifyEnrichment* provides a function `select_cutoff()` which tries a list of cutoffs and compares difference scores, numbers of clusters and block mean values for the clusterings to decide an optimized cutoff value. In [Supplementary file 9](https://simplifyenrichment.github.io/supplementary/suppl9_select_cutoff/suppl9_select_cutoff.html), we demonstrated the use of `select_cutoff()` and how it helps to select a proper cutoff.

Using randomly selected GO terms and a semantic similarity measure, we first compared binary cut with 10 different clustering methods. While some methods were generally characterized by over-segmentation (dynamicTreeCut, mclust and apcluster), others generated intermediate numbers of clusters but either still failed to preserve large clusters with high within-cluster similarities (kmeans) or the largest cluster did not show consistent similarities for all term pairs and should be split further (hdbscan). Even other methods identified both small and large clusters (community methods like fast greedy, leading eigen, louvain, walktrap and MCL), however, even for this latter group, some large clusters may still be split further. As opposed to that, binary cut was able to simultaneously identify clean small and large clusters. Superior performance was also demonstrated quantitatively using various metrics, in particular binary cut had the highest difference score, reflecting the most distinct differences in similarity values between within-clusters and between-clusters comparisons.

It is worth noting that when using semantic measures, even for a list of random GO terms the similarity matrices show clear diagonal block patterns. Semantic measures are based on information content (IC); they rely on the IC of the most informative common ancestor of two terms in the GO tree. A GO term has higher IC if it has less offspring terms. Thus, in general, the deeper the ancestor of two terms is in the GO tree, the higher IC the ancestor will provide and the higher similarity the two terms will have. Uniformly sampling the GO terms selects more terms locating in the downstream of the GO tree due to its hierarchical structure, therefore, it is more probable for two terms to reach an ancestor node deeper in the GO tree to contribute a higher IC value, which infers that even random GO lists can have clear diagonal block patterns in the benchmark.

The performance of *simplifyEnrichment* was also assessed on actual biological data. To this end, we selected datasets from the Expression Atlas based on the number of significantly differentially expressed genes and the number of significant GO terms after functional enrichment with the GO Biological Process ontology. Again, binary cut outperformed other clustering methods. Binary cut also outperformed other clustering methods when applied to random GO lists from the Cellular Component (CC) and Molecular Function (MF) ontologies.

As many ontologies (*e*.*g*. MsigDB gene sets) are only represented as lists of genes for which the similarities are merely measured by various gene overlap metrics while no semantic similarity measure is available, we performed a second benchmark and compared the performance of binary cut when applied to similarity matrices calculated by different similarity measures, again using both randomly selected GO terms and real data from the Expression Atlas. With Jaccard coefficient, Dice coefficient or kappa coefficient as similarity measures, a large number of very small clusters and a small number of bigger clusters were observed on random GO terms, while the overlap coefficient led to massive under-segmentation. As opposed to that, with semantic similarity, these distributions were more equilibrated, intermediate numbers of clusters were generated and the similarity matrices had clear diagonal block patterns. Also on the real-world datasets, the semantic similarity measure worked better on GO terms than gene overlap similarity measures, even though the latter showed more diverse performance. When extending the benchmark to other ontologies, we observed that only gene overlap coefficients of Reactome and MsigDB C4 can be used for clustering with binary cut, while DO, KEGG and other subsets of the MsigDB ontology generally showed low consistency between terms and corresponding similarity matrices did not have clear diagonal block patterns.

Finally, we applied *simplifyEnrichment* to a case study in order to demonstrate its ability to integrate functional enrichment results from multiple lists of genes into one plot. The enrichment results of up-regulated and down-regulated genes in a multi-group differential expression analysis were shown in a compact and intuitive manner. Furthermore, results are fed into word cloud visualization, an additional and unique visualization feature of *simplifyEnrichment*. For even deeper exploration and interactive display of enrichment results, *simplifyEnrichment* has in-built functionality to seamlessly launch a Shiny application with the data in the current workspace of the user.

## CONCLUSIONS

We described a new clustering algorithm, binary cut, for clustering similarity matrices of functional terms. Through comprehensive benchmarks on both simulated and real-world datasets, we demonstrated that binary cut can efficiently cluster functional terms where the terms showed more consistent similarities within clusters and were more mutually exclusive between clusters. We implemented the algorithm into the R package *simplifyEnrichment* which additionally provides functionalities for visualizing, summarizing and comparing the clusterings. We believe *simplifyEnrichment* will be a useful tool for researchers to rapidly explore their results and obtain key biological messages.

## DATA AVAILABILITY

The *simplifyEnrichment* package and the documentation are available at https://bioconductor.org/packages/simplifyEnrichment/. The reports for the analysis of all datasets benchmarked in the paper are available at https://simplifyenrichment.github.io/. The scripts coding the analysis are available at https://github.com/jokergoo/simplifyEnrichment_manuscript. The Expression Atlas datasets are available at https://www.ebi.ac.uk/gxa/download.

## Notes

https://bioconductor.org/packages/simplifyEnrichment/

https://simplifyenrichment.github.io/

### Competing Interest Statement

The authors have declared no competing interest.

### Summary of Updates

Ready for submission.

