## Supplementary file 1 for "*simplifyEnrichment*: an R/Bioconductor package for Clustering and Visualizing Functional Enrichment Results": suppl1_coefficient_definition.html (from another version of this preprint: version 1, 2020-10-28)

Supplementary file 1. Definition of coefficients that measure gene overlaps


### Supplementary file 1. Definition of coefficients that measure gene overlaps

###### Zuguang Gu

#### 2020-09-24

In this supplementary, we give a brief introduction of following coefficients that measure the similarity between two gene sets based on gene overlaps: Jaccard coefficient, Dice coefficient, overlap coefficient and kappa coefficient.

Denote two sets of genes as \(A\) and \(B\), the Jaccard coeffcient between the two sets is calculated as:

\[{|A \cap B|}\over{|A \cup B|}\]

The Dice coeffcient is calculated as:

\[\frac{|A \cap B|}{(|A| + |B|)/2}\]

Jaccard coefficient has a relationship with Dice coefficient in the form of:

\[Jaccard = \frac{Dice}{2 - Dice}\]

The overlap coeffcient is calculated as:

\[\frac{| A \cap B | }{\min(|A|,|B|)}\]

The symbol \(|A|\) means the size of set \(A\) (number of elements).

The definition of kappa coeffient is a little bit complex. First let’s format the two sets into a contigency table:

|  |  |  |  |
| --- | --- | --- | --- |
|  | | In set **B** | |
|  |  | Yes | No |
| In set **A** | Yes | a | b |
|  | No | c | d |

where \(a\), \(b\), \(c\), \(d\) are the numbers of genes that fall in each category.

Let’s calculate \(p\_o\) and \(p\_e\) as:

\[p\_o = \frac{a+d}{a+b+c+d}\]

\[p\_{Yes} = \frac{a+b}{a+b+c+d} \cdot \frac{a+c}{a+b+c+d}\]

\[p\_{No} = \frac{c+d}{a+b+c+d} \cdot \frac{b+d}{a+b+c+d}\]

\[p\_e = p\_{Yes} + p\_{No}\]

where \(p\_o\) is the probability of a gene in both gene sets or neither in the two sets, \(p\_{Yes}\) is the probability of a gene in both gene sets by random (by assuming the events of a gene in set \(A\) and set \(B\) are independent), \(p\_{No}\) is the probability of a gene not in the two sets by random, and \(p\_e\) is the probability of a gene either both in the two sets or not in the two sets by random.

and the kappa coeffcient is calculated as:

\[\frac{p\_o - p\_e}{1 - p\_e}\]
