## Supplementary file 2 for "*simplifyEnrichment*: an R/Bioconductor package for Clustering and Visualizing Functional Enrichment Results": suppl2_random_GO_examples.html (from another version of this preprint: version 1, 2020-10-28)

Supplementary file 2. Semantic similarity matrices of random GO terms


### Supplementary file 2. Semantic similarity matrices of random GO terms

###### Zuguang Gu

#### 2020-08-07

In this supplementary file, we demonstrate the heatmaps of semantic similarities for randomly sampled 500 GO terms, from Biological Process (BP), Molecular Function (MF) and Cellular Component (CC) ontologies. For each ontology category, we show 11 examples.

```
library(simplifyEnrichment)
library(grid)
library(circlize)
library(ComplexHeatmap)
library(GetoptLong)
library(cowplot)

set.seed(123)
col_fun = colorRamp2(c(0, 1), c("white", "red"))
lgd = Legend(title = "Similarity", col_fun = col_fun)
```

#### 500 random BP

```
pl = list()
for(i in 1:11) {
    go_id = random_GO(500, "BP")
    mat = GO_similarity(go_id)
    pl[[i]] = grid.grabExpr(draw(Heatmap(mat, col = col_fun, show_row_names = FALSE, show_column_names = FALSE,
        show_row_dend = FALSE, show_column_dend = FALSE, show_heatmap_legend = FALSE, 
        column_title = qq("random BP #@{i}"))))
}
pl[[12]] = lgd@grob
plot_grid(plotlist = pl, nrow = 3)
```

#### 500 random MF

```
pl = list()
for(i in 1:11) {
    go_id = random_GO(500, "MF")
    mat = GO_similarity(go_id)
    pl[[i]] = grid.grabExpr(draw(Heatmap(mat, col = col_fun, show_row_names = FALSE, show_column_names = FALSE,
        show_row_dend = FALSE, show_column_dend = FALSE, show_heatmap_legend = FALSE, 
        column_title = qq("random MF #@{i}"))))
}
pl[[12]] = lgd@grob
plot_grid(plotlist = pl, nrow = 3)
```

#### 500 random CC

```
pl = list()
for(i in 1:11) {
    go_id = random_GO(500, "CC")
    mat = GO_similarity(go_id)
    pl[[i]] = grid.grabExpr(draw(Heatmap(mat, col = col_fun, show_row_names = FALSE, show_column_names = FALSE,
        show_row_dend = FALSE, show_column_dend = FALSE, show_heatmap_legend = FALSE, 
        column_title = qq("random CC #@{i}"))))
}
pl[[12]] = lgd@grob
plot_grid(plotlist = pl, nrow = 3)
```
