## Supplementary file 3 for "*simplifyEnrichment*: an R/Bioconductor package for Clustering and Visualizing Functional Enrichment Results": suppl3_examples_of_semantic_measures.html (from another version of this preprint: version 1, 2020-10-28)

Supplementary file 3. Examples of semantic similarity measurements


### Supplementary file 3. Examples of semantic similarity measurements

###### Zuguang Gu

#### 2020-10-03

In this supplementary, we demonstrate the heatmaps of semantic similarity matrices from different semantic measurements, which are “Rel”, “Resnik”, “Lin”, “Jiang” and “Wang” (see the GOSemSim vignette for more details of the methods). The matrices are calculated based on 500 randomly sampled GO terms, from Biological Process (BP), Molecular Function (MF) and Cellular Component (CC) ontologies. For each ontology category, we show 5 examples.

Generally, the similarity matrices generated by the “Rel”, “Resnik” and “Lin” methods show more clear diagonal patterns.

#### 500 random BP

##### Random 1

##### Random 2

##### Random 3

##### Random 4

##### Random 5

#### 500 random MF

##### Random 1

##### Random 2

##### Random 3

##### Random 4

##### Random 5

#### 500 random CC

##### Random 1

##### Random 2

##### Random 3

##### Random 4

##### Random 5
