## Supplementary file 4 for "*simplifyEnrichment*: an R/Bioconductor package for Clustering and Visualizing Functional Enrichment Results": suppl4_examples.html (from another version of this preprint: version 1, 2020-10-28)

Supplementary file 4. Examples of simplifyEnrichment


### Supplementary file 4. Examples of simplifyEnrichment

This page contains results for functional enrichment clustering done by the **simplifyEnrichment** package.

---

#### Random GO

- random\_BP (Based on semantic similarity matrix)
- random\_CC (Based on semantic similarity matrix)
- random\_MF (Based on semantic similarity matrix)
- random\_BP\_dice
- random\_BP\_jaccard
- random\_BP\_kappa
- random\_BP\_overlap

---

#### On EBI Expression Atlas datasets

### GO

- EBI\_Expression\_Atlas\_GO\_BP (Based on semantic similarity matrix)
- EBI\_Expression\_Atlas\_GO\_BP\_dice
- EBI\_Expression\_Atlas\_GO\_BP\_jaccard
- EBI\_Expression\_Atlas\_GO\_BP\_kappa
- EBI\_Expression\_Atlas\_GO\_BP\_overlap

### DO

- EBI\_Expression\_Atlas\_DO (Based on semantic similarity matrix)
- EBI\_Expression\_Atlas\_DO\_dice
- EBI\_Expression\_Atlas\_DO\_jaccard
- EBI\_Expression\_Atlas\_DO\_kappa
- EBI\_Expression\_Atlas\_DO\_overlap

##### KEGG

- EBI\_Expression\_Atlas\_KEGG\_dice
- EBI\_Expression\_Atlas\_KEGG\_jaccard
- EBI\_Expression\_Atlas\_KEGG\_kappa
- EBI\_Expression\_Atlas\_KEGG\_overlap

##### Reactome

- EBI\_Expression\_Atlas\_Reactome\_dice
- EBI\_Expression\_Atlas\_Reactome\_jaccard
- EBI\_Expression\_Atlas\_Reactome\_kappa
- EBI\_Expression\_Atlas\_Reactome\_overlap

##### MsigDB\_C2\_CGP

- EBI\_Expression\_Atlas\_MsigDB\_C2\_CGP\_dice
- EBI\_Expression\_Atlas\_MsigDB\_C2\_CGP\_jaccard
- EBI\_Expression\_Atlas\_MsigDB\_C2\_CGP\_kappa
- EBI\_Expression\_Atlas\_MsigDB\_C2\_CGP\_overlap

##### MsigDB\_C3\_GTRD

- EBI\_Expression\_Atlas\_MsigDB\_C3\_GTRD\_dice
- EBI\_Expression\_Atlas\_MsigDB\_C3\_GTRD\_jaccard
- EBI\_Expression\_Atlas\_MsigDB\_C3\_GTRD\_kappa
- EBI\_Expression\_Atlas\_MsigDB\_C3\_GTRD\_overlap

##### MsigDB\_C3\_MIR\_Legacy

- EBI\_Expression\_Atlas\_MsigDB\_C3\_MIR\_Legacy\_dice
- EBI\_Expression\_Atlas\_MsigDB\_C3\_MIR\_Legacy\_jaccard
- EBI\_Expression\_Atlas\_MsigDB\_C3\_MIR\_Legacy\_kappa
- EBI\_Expression\_Atlas\_MsigDB\_C3\_MIR\_Legacy\_overlap

##### MsigDB\_C3\_MIRDB

- EBI\_Expression\_Atlas\_MsigDB\_C3\_MIRDB\_dice
- EBI\_Expression\_Atlas\_MsigDB\_C3\_MIRDB\_jaccard
- EBI\_Expression\_Atlas\_MsigDB\_C3\_MIRDB\_kappa
- EBI\_Expression\_Atlas\_MsigDB\_C3\_MIRDB\_overlap

##### MsigDB\_C3\_TFT\_Legacy

- EBI\_Expression\_Atlas\_MsigDB\_C3\_TFT\_Legacy\_dice
- EBI\_Expression\_Atlas\_MsigDB\_C3\_TFT\_Legacy\_jaccard
- EBI\_Expression\_Atlas\_MsigDB\_C3\_TFT\_Legacy\_kappa
- EBI\_Expression\_Atlas\_MsigDB\_C3\_TFT\_Legacy\_overlap

##### MsigDB\_C4\_CGN

- EBI\_Expression\_Atlas\_MsigDB\_C4\_CGN\_dice
- EBI\_Expression\_Atlas\_MsigDB\_C4\_CGN\_jaccard
- EBI\_Expression\_Atlas\_MsigDB\_C4\_CGN\_kappa
- EBI\_Expression\_Atlas\_MsigDB\_C4\_CGN\_overlap

##### MsigDB\_C4\_CM

- EBI\_Expression\_Atlas\_MsigDB\_C4\_CM\_dice
- EBI\_Expression\_Atlas\_MsigDB\_C4\_CM\_jaccard
- EBI\_Expression\_Atlas\_MsigDB\_C4\_CM\_kappa
- EBI\_Expression\_Atlas\_MsigDB\_C4\_CM\_overlap

##### MsigDB\_C7

- EBI\_Expression\_Atlas\_MsigDB\_C7\_\_dice
- EBI\_Expression\_Atlas\_MsigDB\_C7\_\_jaccard
- EBI\_Expression\_Atlas\_MsigDB\_C7\_\_kappa
- EBI\_Expression\_Atlas\_MsigDB\_C7\_\_overlap
