## Supplementary file 5 for "*simplifyEnrichment*: an R/Bioconductor package for Clustering and Visualizing Functional Enrichment Results": suppl5_compare_similarity_random_GO.html (from another version of this preprint: version 1, 2020-10-28)

Supplementary 5. Compare similarity measurements - random\_BP


### Supplementary 5. Compare similarity measurements - random\_BP

---

**Figure S5.1.** Left) Numbers of clusters. Middle) Average numbers of terms per cluster. Right) Concordance of clusterings from different similarity matrices. The definition of the concordance score can be found here.

  

**Figure S5.2.**Numbers of clusters.

  

**Figure S5.3.**Numbers of large clusters (size >= 5).

  

**Figure S5.4.**Proportion of the largest cluster.

  

**Table S5.1.** Details on individual datasets.

|  |  |  |  |  |  |  |  |  |  |
| --- | --- | --- | --- | --- | --- | --- | --- | --- | --- |
| random\_BP\_1 | random\_BP\_2 | random\_BP\_3 | random\_BP\_4 | random\_BP\_5 | random\_BP\_6 | random\_BP\_7 | random\_BP\_8 | random\_BP\_9 | random\_BP\_10 |
| random\_BP\_11 | random\_BP\_12 | random\_BP\_13 | random\_BP\_14 | random\_BP\_15 | random\_BP\_16 | random\_BP\_17 | random\_BP\_18 | random\_BP\_19 | random\_BP\_20 |
| random\_BP\_21 | random\_BP\_22 | random\_BP\_23 | random\_BP\_24 | random\_BP\_25 | random\_BP\_26 | random\_BP\_27 | random\_BP\_28 | random\_BP\_29 | random\_BP\_30 |
| random\_BP\_31 | random\_BP\_32 | random\_BP\_33 | random\_BP\_34 | random\_BP\_35 | random\_BP\_36 | random\_BP\_37 | random\_BP\_38 | random\_BP\_39 | random\_BP\_40 |
| random\_BP\_41 | random\_BP\_42 | random\_BP\_43 | random\_BP\_44 | random\_BP\_45 | random\_BP\_46 | random\_BP\_47 | random\_BP\_48 | random\_BP\_49 | random\_BP\_50 |
| random\_BP\_51 | random\_BP\_52 | random\_BP\_53 | random\_BP\_54 | random\_BP\_55 | random\_BP\_56 | random\_BP\_57 | random\_BP\_58 | random\_BP\_59 | random\_BP\_60 |
| random\_BP\_61 | random\_BP\_62 | random\_BP\_63 | random\_BP\_64 | random\_BP\_65 | random\_BP\_66 | random\_BP\_67 | random\_BP\_68 | random\_BP\_69 | random\_BP\_70 |
| random\_BP\_71 | random\_BP\_72 | random\_BP\_73 | random\_BP\_74 | random\_BP\_75 | random\_BP\_76 | random\_BP\_77 | random\_BP\_78 | random\_BP\_79 | random\_BP\_80 |
| random\_BP\_81 | random\_BP\_82 | random\_BP\_83 | random\_BP\_84 | random\_BP\_85 | random\_BP\_86 | random\_BP\_87 | random\_BP\_88 | random\_BP\_89 | random\_BP\_90 |
| random\_BP\_91 | random\_BP\_92 | random\_BP\_93 | random\_BP\_94 | random\_BP\_95 | random\_BP\_96 | random\_BP\_97 | random\_BP\_98 | random\_BP\_99 | random\_BP\_100 |
