## Supplementary figures and images for "*simplifyEnrichment*: an R/Bioconductor package for Clustering and Visualizing Functional Enrichment Results" (from another version of this preprint: version 1, 2020-10-28)

### random_BP_compare_similarity.png

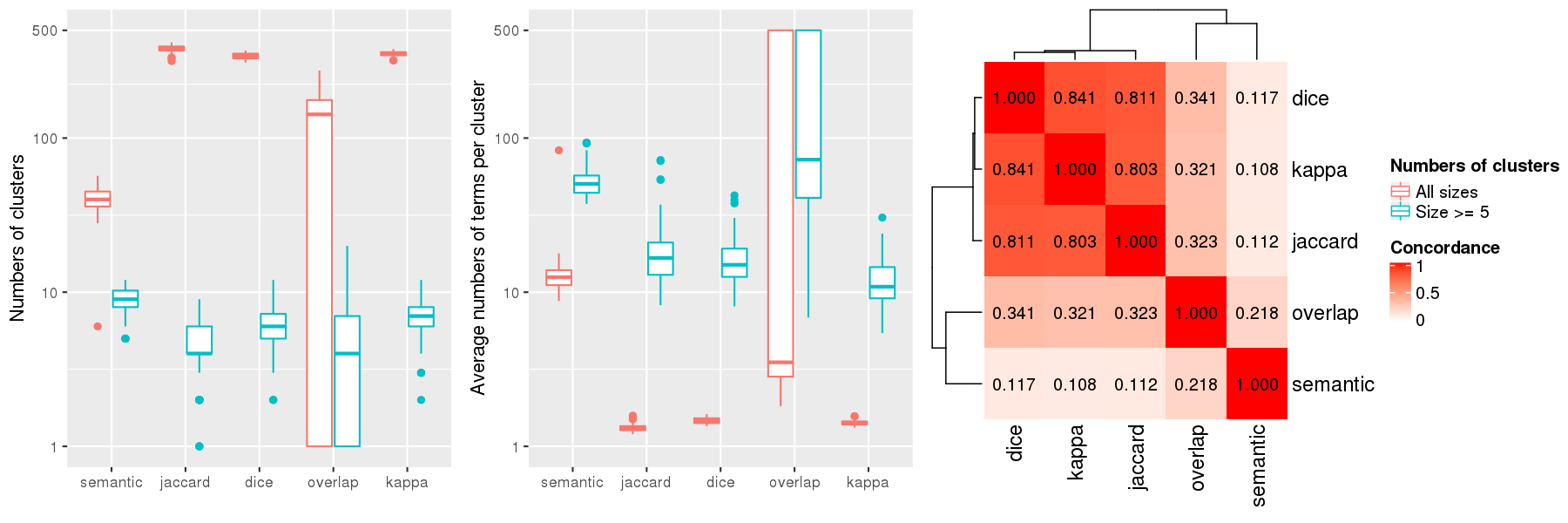

### random_BP_compare_similarity_large_ncluster_barplot.png

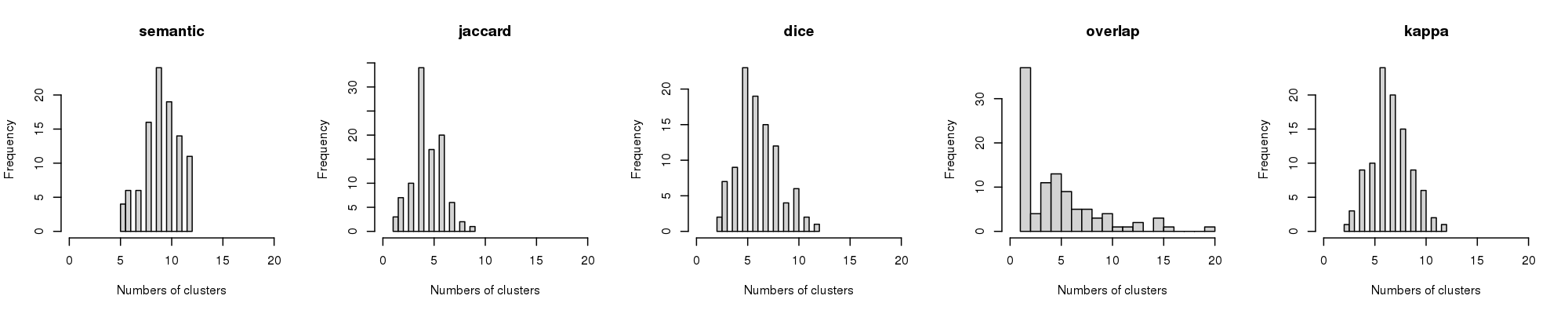

### random_BP_compare_similarity_max_cluster_prop_barplot.png

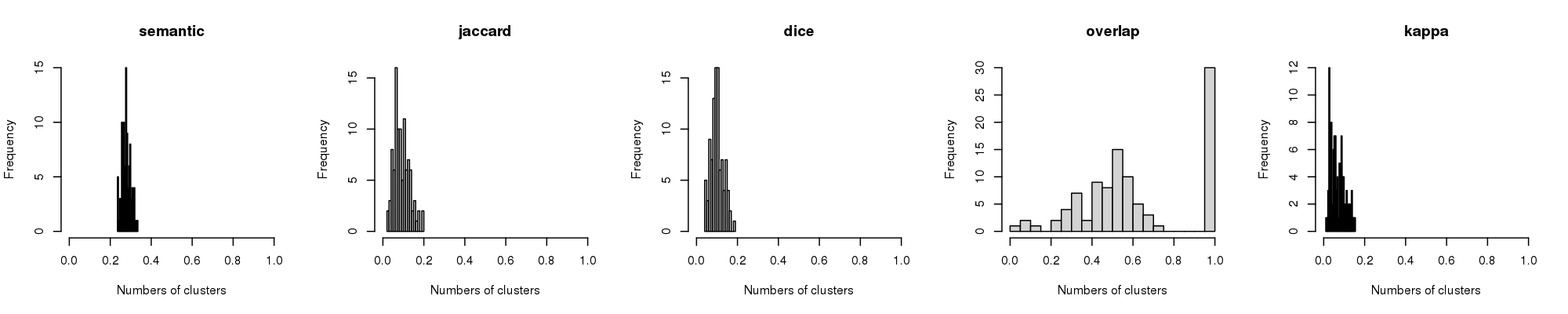

### random_BP_compare_similarity_ncluster_barplot.png

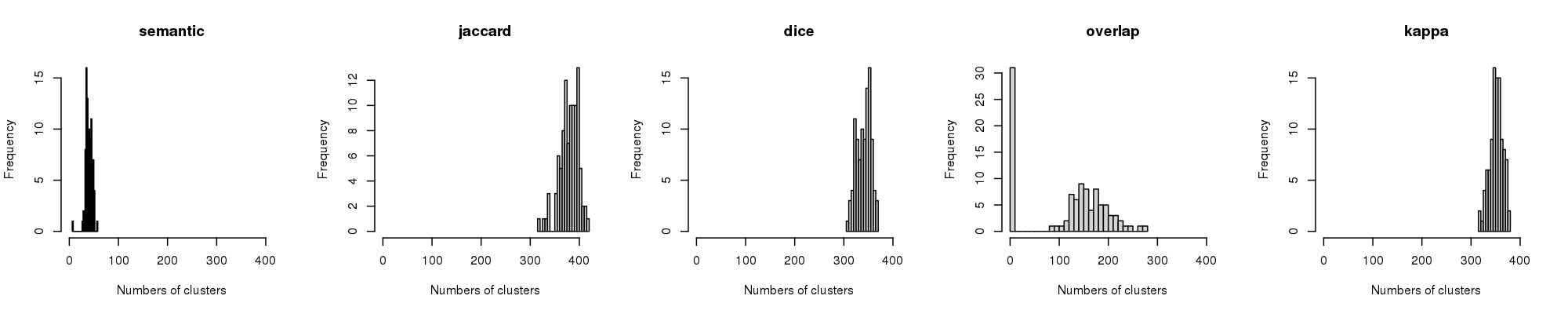
