## Supplementary file 6 for "*simplifyEnrichment*: an R/Bioconductor package for Clustering and Visualizing Functional Enrichment Results": suppl6_compare_similarity.html (from another version of this preprint: version 1, 2020-10-28)

Supplementary file 6. Compare similarity measurements


### Supplementary file 6. Compare similarity measurements

---

We compared the clusterings on the similarity matrices with different similarity measurements.

| dataset | semantic | Jaccard | Dice | overlap | kappa |
| --- | --- | --- | --- | --- | --- |
| Compare similarity measurements - random\_GO\_BP | x | x | x | x | x |
| Compare similarity measurements - EBI\_Expression\_Atlas\_GO\_BP | x | x | x | x | x |
| Compare similarity measurements - EBI\_Expression\_Atlas\_DO | x | x | x | x | x |
| Compare similarity measurements - EBI\_Expression\_Atlas\_KEGG |  | x | x | x | x |
| Compare similarity measurements - EBI\_Expression\_Atlas\_Reactome |  | x | x | x | x |
| Compare similarity measurements - EBI\_Expression\_Atlas\_MsigDB\_C2\_CGP |  | x | x | x | x |
| Compare similarity measurements - EBI\_Expression\_Atlas\_MsigDB\_C3\_GTRD |  | x | x | x | x |
| Compare similarity measurements - EBI\_Expression\_Atlas\_MsigDB\_C3\_MIR\_Legacy |  | x | x | x | x |
| Compare similarity measurements - EBI\_Expression\_Atlas\_MsigDB\_C3\_MIRDB |  | x | x | x | x |
| Compare similarity measurements - EBI\_Expression\_Atlas\_MsigDB\_C3\_TFT\_Legacy |  | x | x | x | x |
| Compare similarity measurements - EBI\_Expression\_Atlas\_MsigDB\_C4\_CGN |  | x | x | x | x |
| Compare similarity measurements - EBI\_Expression\_Atlas\_MsigDB\_C4\_CM |  | x | x | x | x |
| Compare similarity measurements - EBI\_Expression\_Atlas\_MsigDB\_C7 |  | x | x | x | x |
