## Supplementary file 7 for "*simplifyEnrichment*: an R/Bioconductor package for Clustering and Visualizing Functional Enrichment Results": suppl7_global_GO_similarity.html (from another version of this preprint: version 1, 2020-10-28)

Supplementary file 7. Global GO term similarity


### Supplementary file 7. Global GO term similarity

###### Zuguang Gu

#### 2020-09-20

In this supplementary, we look at the global GO term similarities for Biological Process (BP) ontology. First we calculate the similarity matrix for all GO BP terms supported in *GOSemSim* package.

```
library(GOSemSim)
semData = godata("org.Hs.eg.db", ont = "BP")
ic = semData@IC

library(simplifyEnrichment)
# This takes very long time
sm = GO_similarity(names(ic), measure = "Rel", ont = "BP")
```

The number of all GO terms in BP ontology:

```
dim(sm)
```

```
## [1] 27918 27918
```

To simplify the similarity matrix, the GO terms which locate as leaves in the GO tree are removed. These leaf terms only have similarity with themselves, thus they have zero similarity to all other terms.

```
l = rowSums(sm > 0) == 1
sm = sm[!l, !l]
```

Next we construct the GO tree. The parent-children information is from *GO.db* package. The `GOBPCHILDREN` data object contains lists of direct children terms for every GO terms. If a GO term is a leaf in the GO tree, the value for it is `NA`.

```
library(GO.db)
lt = as.list(GOBPCHILDREN)
lt[1:5]
```

```
## $`GO:0000001`
## [1] NA
## 
## $`GO:0000002`
##      part of          isa 
## "GO:0032042" "GO:0033955" 
## 
## $`GO:0000003`
##          isa          isa      part of          isa          isa          isa          isa 
## "GO:0019953" "GO:0019954" "GO:0022414" "GO:0032504" "GO:0032505" "GO:0061887" "GO:0075325" 
## 
## $`GO:0000011`
## [1] NA
## 
## $`GO:0000012`
##            regulates negatively regulates positively regulates                  isa 
##         "GO:1903516"         "GO:1903517"         "GO:1903518"         "GO:1903823" 
##                  isa 
##         "GO:1990396"
```

Then we can construct a graph object for the GO tree.

```
df = do.call("rbind", lapply(names(lt), function(nm) {
    data.frame(rep(nm, length(lt[[nm]])), lt[[nm]])
}))
df2 = df[!is.na(df[, 2]), ]

library(igraph)
g = graph.edgelist(as.matrix(df2), directed = TRUE)
g
```

```
## IGRAPH d34cbf9 DN-- 29211 70659 -- 
## + attr: name (v/c)
## + edges from d34cbf9 (vertex names):
##  [1] GO:0000002->GO:0032042 GO:0000002->GO:0033955 GO:0000003->GO:0019953 GO:0000003->GO:0019954
##  [5] GO:0000003->GO:0022414 GO:0000003->GO:0032504 GO:0000003->GO:0032505 GO:0000003->GO:0061887
##  [9] GO:0000003->GO:0075325 GO:0000012->GO:1903516 GO:0000012->GO:1903517 GO:0000012->GO:1903518
## [13] GO:0000012->GO:1903823 GO:0000012->GO:1990396 GO:0000018->GO:0000019 GO:0000018->GO:0000337
## [17] GO:0000018->GO:0010520 GO:0000018->GO:0010569 GO:0000018->GO:0045191 GO:0000018->GO:0045910
## [21] GO:0000018->GO:0045911 GO:0000018->GO:0060542 GO:0000018->GO:0061806 GO:0000018->GO:0072695
## [25] GO:0000018->GO:1903110 GO:0000019->GO:0032207 GO:0000019->GO:0045950 GO:0000019->GO:0045951
## [29] GO:0000019->GO:1903221 GO:0000022->GO:0032888 GO:0000022->GO:0051256 GO:0000022->GO:0061804
## + ... omitted several edges
```

We take the common GO terms in the GO tree and the global similarity matrix.

```
cn = intersect(rownames(sm), unlist(df2))
sm2 = sm[cn, cn]
g = induced_subgraph(g, cn)
sm2 = sm2[V(g)$name, V(g)$name]
```

The final number of GO terms are:

```
dim(sm2)
```

```
## [1] 15271 15271
```

The gobal similarity matrix is clustered by the “louvain” method. Note binary cut is too slow to work on a similarity matrix with 15k GO terms.

```
# This takes around 30 min
cl = cluster_terms(sm2, method = "louvain")
```

The size of clusters:

```
table(cl)
```

```
## cl
##    1    2    3    4    5 
## 2332 3996 4329 1074 3540
```

To visualize the similarity matrix and the clustering, we only randomly sample 2000 terms.

```
set.seed(123)
ind = sample(length(cl), 2000)
ht_clusters(sm2[ind, ind], cl = cl[ind], exclude_words = c("process", "regulation", "response"))
```

Next we look at what are the top GO terms in each cluster (*i.e.*, the GO terms located on the top of the GO subtree in each cluster). First we construct subgraph for each cluster.

```
gl = tapply(rownames(sm2), cl, function(go_id) {
    induced_subgraph(g, go_id)
})
```

Since the subgraphs are split from the global tree/graph, they might have several disconnected compoments. We check the sizes of disconnected components in each subgraph.

```
lapply(gl, function(g) {
    table(components(g)$membership)
})
```

```
## $`1`
## 
##    1    2    3    4    5    6    7    8    9   10   11 
## 2320    1    1    3    1    1    1    1    1    1    1 
## 
## $`2`
## 
##    1    2    3    4    5    6 
## 3991    1    1    1    1    1 
## 
## $`3`
## 
##    1    2    3    4    5    6    7    8    9   10   11   12   13   14 
## 4311    3    1    1    1    2    1    1    1    3    1    1    1    1 
## 
## $`4`
## 
##    1    2    3    4    5    6    7    8    9   10   11   12   13   14   15 
## 1031    3   10    6    5    3    1    4    2    4    1    1    1    1    1 
## 
## $`5`
## 
##    1    2    3    4    5    6    7    8    9   10   11 
## 3522    3    1    1    1    1    1    1    1    1    7
```

As we can see, in each cluster, there is always a huge components which have majority of the terms while other components have very few terms. Thus, we only take the componets with the largest number of terms.

```
gl2 = lapply(gl, function(g) {
    membership = components(g)$membership
    induced_subgraph(g, membership == which.max(table(membership)))
})
```

Next we extract the top GO terms where the in-degree is zero. And we only want the GO terms which are the parent nodes of a huge GO subtree. The influences a parent node gives to all its descendant nodes is measured by the “spread centrality”.

```
library(CePa)
lapply(gl2, function(g) {
    s1 = degree(g, mode = "in")
    s2 = spread(g, mode = "out")
    s2[s1 == 0]
})
```

```
## $`1`
## GO:0006928 GO:0008089 GO:0008090 GO:0040011 GO:0051179 GO:0044417 GO:0065008 GO:0071539 GO:0071988 
##   56.81429    4.50000    1.00000   37.04762  463.42421    2.50000  147.99293    3.00000    2.00000 
## GO:0090670 
##    3.00000 
## 
## $`2`
## GO:0000003 GO:0003176 GO:0003177 GO:0007610 GO:0032501 GO:0032502 GO:0040007 GO:0044848 GO:0048511 
##   73.55238    1.00000    1.50000   35.51667  860.38280  930.24325   52.67857    4.00000   18.11667 
## GO:0098743 GO:0042310 GO:0042311 GO:0072673 
##    1.00000    8.50000    4.00000    3.00000 
## 
## $`3`
## GO:0000045 GO:0008152 GO:0009987 GO:0010412 GO:0061724 
##     5.0000  1002.9036   778.3294     1.0000     2.0000 
## 
## $`4`
## GO:0007017 GO:0007049 GO:0007059 GO:0007127 GO:0007135 GO:0007140 GO:0007143 GO:0007163 GO:0030029 
##  34.483333  90.783333  29.792857   9.083333   3.500000   3.500000   7.000000  10.333333  22.361905 
## GO:0033059 GO:0045103 GO:0051301 GO:0071840 GO:2000241 GO:0032528 GO:0097061 GO:0097581 GO:1904868 
##   2.666667   4.250000  15.833333 214.739683   7.650000   3.500000   2.500000   5.500000   2.000000 
## 
## $`5`
## GO:0001775 GO:0001906 GO:0002376 GO:0007154 GO:0007268 GO:0009758 GO:0019740 GO:0022610 GO:0023052 
##  62.442857  28.733333 178.847619 252.498810  33.116667   1.000000   4.000000  36.619048 238.582143 
## GO:0044419 GO:0050896 GO:0051703 GO:0051704 GO:0065007 GO:0008219 GO:0008283 GO:0019835 GO:0016055 
##  98.552381 619.622222   4.166667  11.266667 423.985714  78.911905  82.319048   7.833333  16.000000 
## GO:0042710 
##   3.166667
```

As shown above, in each cluster, there are some top GO terms that are only the parents of a small number of children terms while some other terms have extremely high spread values which means they are parents of huge number of children terms. We simply remove the top GO terms with spread values less than the mean spread values of all top GO terms.

```
go_list = lapply(gl2, function(g) {
    s1 = degree(g, mode = "in")
    s2 = spread(g, mode = "out")
    go_id = V(g)$name[which(s1 == 0)]
    s = s2[s1 == 0]
    go_id = go_id[s >= mean(s)]
    term = suppressMessages(select(GO.db::GO.db, keys = go_id, columns = "TERM")$TERM)
    paste0(go_id, ": ", term)
})
```

The final major top GO terms in each cluster are:

```
go_list
```

```
## $`1`
## [1] "GO:0051179: localization"                    
## [2] "GO:0065008: regulation of biological quality"
## 
## $`2`
## [1] "GO:0032501: multicellular organismal process"
## [2] "GO:0032502: developmental process"           
## 
## $`3`
## [1] "GO:0008152: metabolic process"
## [2] "GO:0009987: cellular process" 
## 
## $`4`
## [1] "GO:0007017: microtubule-based process"                    
## [2] "GO:0007049: cell cycle"                                   
## [3] "GO:0007059: chromosome segregation"                       
## [4] "GO:0071840: cellular component organization or biogenesis"
## 
## $`5`
## [1] "GO:0002376: immune system process"
## [2] "GO:0007154: cell communication"   
## [3] "GO:0023052: signaling"            
## [4] "GO:0050896: response to stimulus" 
## [5] "GO:0065007: biological regulation"
```
