## Supplementary file 8 for "*simplifyEnrichment*: an R/Bioconductor package for Clustering and Visualizing Functional Enrichment Results": suppl8_partitioning_methods.html (from another version of this preprint: version 1, 2020-10-28)

Supplementary file 8. Test partitioning methods


### Supplementary file 8. Test partitioning methods

---

In the process of recursively splitting the similarity matrix in binary cut algorithm,
in each iteration step, the current matrix is partitioned into two groups using PAM as default.
Here we compare following partitioning methds: k-means, PAM and hierarchical clustering with
methods of 'complete', 'average' and 'ward.D2', on 500 random GO lists.

**Figure S8.1.**Compare clustering results. Left panel: The difference score, number of clusters and the block mean of different clusterings. Right panel: Concordance between clustering methods. The concordance measures how similar two clusterings are. The definition of the concordance score can be found here.

**Table S8.1.**Number of clusters identified by each clustering method. Numbers in the table indicate the number of clusters. The numbers inside the parentheses are the number of clusters with size >= 5.

| run | partition\_by\_kmeans | partition\_by\_pam | partition\_by\_hclust\_complete | partition\_by\_hclust\_ward.D2 | partition\_by\_hclust\_average | Details |
| --- | --- | --- | --- | --- | --- | --- |
| 1 | 47 (9) | 38 (7) | 41 (12) | 34 (6) | 37 (7) | view |
| 2 | 32 (9) | 27 (5) | 32 (8) | 31 (10) | 32 (9) | view |
| 3 | 43 (10) | 42 (12) | 46 (13) | 41 (12) | 44 (13) | view |
| 4 | 41 (9) | 38 (6) | 40 (14) | 38 (10) | 37 (12) | view |
| 5 | 42 (12) | 36 (11) | 35 (11) | 30 (9) | 33 (10) | view |
| 6 | 38 (7) | 40 (10) | 38 (10) | 36 (9) | 38 (10) | view |
| 7 | 31 (10) | 31 (11) | 31 (11) | 27 (8) | 27 (10) | view |
| 8 | 38 (9) | 36 (9) | 35 (9) | 35 (9) | 35 (12) | view |
| 9 | 46 (9) | 39 (11) | 40 (13) | 39 (11) | 38 (11) | view |
| 10 | 37 (9) | 37 (8) | 40 (13) | 35 (8) | 38 (8) | view |
| 11 | 52 (12) | 41 (9) | 39 (15) | 36 (9) | 38 (10) | view |
| 12 | 43 (9) | 41 (11) | 40 (10) | 40 (9) | 39 (9) | view |
| 13 | 34 (12) | 30 (10) | 33 (10) | 30 (8) | 33 (11) | view |
| 14 | 45 (8) | 36 (9) | 35 (10) | 33 (9) | 34 (9) | view |
| 15 | 49 (8) | 42 (10) | 41 (12) | 38 (10) | 40 (9) | view |
| 16 | 45 (9) | 36 (12) | 36 (12) | 33 (12) | 34 (12) | view |
| 17 | 49 (16) | 44 (13) | 45 (13) | 42 (12) | 43 (13) | view |
| 18 | 39 (10) | 40 (9) | 41 (14) | 36 (10) | 43 (15) | view |
| 19 | 40 (9) | 40 (10) | 42 (14) | 35 (8) | 37 (9) | view |
| 20 | 44 (9) | 38 (9) | 39 (9) | 37 (8) | 41 (8) | view |
| 21 | 48 (12) | 40 (10) | 40 (12) | 38 (10) | 38 (9) | view |
| 22 | 36 (10) | 37 (11) | 34 (9) | 35 (12) | 36 (12) | view |
| 23 | 37 (7) | 33 (8) | 35 (10) | 31 (9) | 32 (10) | view |
| 24 | 44 (11) | 40 (10) | 35 (12) | 34 (10) | 37 (11) | view |
| 25 | 46 (8) | 42 (9) | 43 (11) | 40 (11) | 43 (13) | view |
| 26 | 43 (9) | 41 (9) | 47 (13) | 38 (5) | 43 (10) | view |
| 27 | 39 (11) | 38 (10) | 39 (11) | 37 (10) | 38 (10) | view |
| 28 | 52 (15) | 37 (13) | 41 (16) | 37 (13) | 37 (14) | view |
| 29 | 38 (10) | 37 (13) | 39 (11) | 33 (9) | 37 (12) | view |
| 30 | 36 (6) | 35 (8) | 33 (8) | 32 (7) | 32 (7) | view |
| 31 | 47 (9) | 41 (11) | 42 (14) | 36 (9) | 38 (9) | view |
| 32 | 38 (8) | 42 (9) | 40 (11) | 36 (7) | 40 (8) | view |
| 33 | 37 (8) | 38 (9) | 37 (11) | 36 (10) | 36 (10) | view |
| 34 | 39 (11) | 34 (10) | 28 (9) | 27 (9) | 31 (9) | view |
| 35 | 38 (11) | 38 (9) | 38 (9) | 32 (7) | 38 (9) | view |
| 36 | 38 (11) | 33 (9) | 32 (9) | 32 (8) | 32 (9) | view |
| 37 | 52 (15) | 48 (11) | 48 (13) | 45 (11) | 47 (11) | view |
| 38 | 32 (11) | 28 (16) | 32 (17) | 25 (12) | 24 (12) | view |
| 39 | 42 (12) | 41 (12) | 40 (11) | 37 (12) | 41 (12) | view |
| 40 | 44 (6) | 39 (9) | 41 (12) | 36 (8) | 38 (10) | view |
| 41 | 37 (8) | 36 (9) | 36 (10) | 34 (9) | 0 (0) | view |
| 42 | 28 (7) | 33 (12) | 28 (9) | 28 (7) | 29 (9) | view |
| 43 | 41 (13) | 34 (10) | 35 (9) | 35 (9) | 37 (10) | view |
| 44 | 35 (13) | 31 (10) | 36 (11) | 31 (9) | 30 (9) | view |
| 45 | 48 (10) | 42 (9) | 43 (9) | 39 (7) | 41 (7) | view |
| 46 | 40 (7) | 38 (8) | 38 (10) | 37 (7) | 39 (9) | view |
| 47 | 44 (10) | 37 (8) | 37 (9) | 35 (9) | 37 (10) | view |
| 48 | 34 (10) | 31 (7) | 32 (10) | 31 (8) | 31 (10) | view |
| 49 | 37 (10) | 29 (9) | 32 (10) | 33 (10) | 33 (9) | view |
| 50 | 37 (9) | 39 (10) | 38 (10) | 34 (9) | 37 (10) | view |
| 51 | 42 (8) | 37 (10) | 41 (12) | 38 (10) | 39 (10) | view |
| 52 | 35 (11) | 30 (11) | 28 (11) | 28 (10) | 30 (10) | view |
| 53 | 45 (8) | 43 (8) | 41 (10) | 41 (9) | 40 (10) | view |
| 54 | 41 (7) | 39 (10) | 42 (13) | 38 (9) | 37 (9) | view |
| 55 | 50 (15) | 37 (14) | 36 (14) | 33 (12) | 35 (13) | view |
| 56 | 46 (11) | 36 (9) | 38 (14) | 35 (8) | 35 (10) | view |
| 57 | 42 (8) | 35 (10) | 35 (11) | 33 (9) | 31 (8) | view |
| 58 | 30 (9) | 29 (10) | 31 (10) | 29 (10) | 29 (9) | view |
| 59 | 37 (12) | 34 (11) | 34 (9) | 35 (8) | 34 (10) | view |
| 60 | 37 (9) | 35 (8) | 36 (10) | 35 (7) | 37 (8) | view |
| 61 | 46 (11) | 40 (11) | 39 (12) | 39 (13) | 39 (12) | view |
| 62 | 38 (11) | 35 (11) | 34 (10) | 35 (10) | 38 (11) | view |
| 63 | 38 (14) | 34 (10) | 32 (8) | 31 (8) | 33 (8) | view |
| 64 | 46 (9) | 41 (9) | 41 (9) | 39 (8) | 40 (8) | view |
| 65 | 42 (10) | 36 (6) | 42 (12) | 36 (9) | 36 (9) | view |
| 66 | 39 (9) | 35 (7) | 37 (10) | 33 (9) | 0 (0) | view |
| 67 | 39 (9) | 33 (10) | 37 (11) | 32 (9) | 34 (10) | view |
| 68 | 45 (9) | 38 (11) | 36 (10) | 36 (10) | 37 (10) | view |
| 69 | 39 (9) | 31 (11) | 31 (11) | 30 (9) | 28 (9) | view |
| 70 | 35 (8) | 32 (11) | 33 (11) | 30 (9) | 32 (11) | view |
| 71 | 22 (6) | 32 (10) | 33 (9) | 34 (10) | 35 (11) | view |
| 72 | 35 (10) | 34 (10) | 33 (12) | 33 (10) | 34 (10) | view |
| 73 | 54 (11) | 42 (8) | 44 (13) | 39 (10) | 39 (9) | view |
| 74 | 43 (15) | 34 (12) | 35 (12) | 34 (10) | 36 (11) | view |
| 75 | 36 (9) | 36 (8) | 37 (14) | 33 (11) | 33 (9) | view |
| 76 | 47 (7) | 44 (7) | 43 (9) | 43 (8) | 43 (9) | view |
| 77 | 39 (7) | 39 (9) | 41 (11) | 41 (11) | 43 (10) | view |
| 78 | 36 (10) | 32 (8) | 30 (8) | 30 (9) | 32 (8) | view |
| 79 | 44 (10) | 39 (8) | 38 (8) | 37 (7) | 38 (9) | view |
| 80 | 39 (9) | 34 (6) | 42 (12) | 40 (12) | 39 (10) | view |
| 81 | 47 (11) | 47 (11) | 47 (12) | 43 (8) | 47 (12) | view |
| 82 | 46 (13) | 42 (12) | 40 (10) | 41 (10) | 41 (10) | view |
| 83 | 33 (8) | 32 (9) | 34 (8) | 30 (8) | 33 (9) | view |
| 84 | 37 (10) | 36 (10) | 36 (9) | 35 (9) | 35 (10) | view |
| 85 | 43 (9) | 41 (11) | 37 (13) | 34 (8) | 35 (10) | view |
| 86 | 36 (8) | 34 (10) | 35 (13) | 30 (9) | 34 (10) | view |
| 87 | 39 (8) | 38 (10) | 37 (11) | 35 (9) | 38 (11) | view |
| 88 | 29 (6) | 32 (10) | 32 (12) | 30 (9) | 29 (10) | view |
| 89 | 37 (9) | 34 (9) | 35 (11) | 33 (10) | 33 (9) | view |
| 90 | 48 (11) | 42 (11) | 39 (8) | 42 (10) | 39 (8) | view |
| 91 | 40 (8) | 41 (9) | 38 (9) | 39 (10) | 38 (9) | view |
| 92 | 31 (11) | 31 (11) | 32 (12) | 29 (8) | 31 (10) | view |
| 93 | 36 (10) | 35 (9) | 36 (10) | 34 (9) | 35 (9) | view |
| 94 | 36 (12) | 36 (13) | 37 (13) | 35 (12) | 36 (10) | view |
| 95 | 44 (12) | 34 (9) | 41 (15) | 39 (12) | 39 (12) | view |
| 96 | 47 (13) | 40 (10) | 42 (11) | 37 (6) | 40 (8) | view |
| 97 | 45 (9) | 35 (10) | 38 (12) | 34 (9) | 34 (9) | view |
| 98 | 38 (10) | 32 (11) | 31 (11) | 30 (11) | 33 (11) | view |
| 99 | 33 (9) | 36 (9) | 37 (10) | 33 (6) | 32 (8) | view |
| 100 | 39 (9) | 40 (10) | 41 (12) | 42 (11) | 40 (10) | view |
