## Supplementary file 9 for "*simplifyEnrichment*: an R/Bioconductor package for Clustering and Visualizing Functional Enrichment Results": suppl9_select_cutoff.html (from another version of this preprint: version 1, 2020-10-28)

Supplementary file 9. Select cutoff in binary cut clustering


### Supplementary file 9. Select cutoff in binary cut clustering

###### Zuguang Gu

#### 2020-09-21

In every step of binary cut clustering, a score is calculated based on the two-group partitioning and it is compared against a cutoff to decide whether the current matrix is treated as a single group or should be split more. In this document, we demonstrate how to fine-tune the cutoff to get nicer clusterings.

```
library(simplifyEnrichment)
library(grid)
```

The default cutoff is 0.85, it works well if the diagonal pattern is obvious from the heatmap.

```
rl = readRDS("/icgc/dkfzlsdf/analysis/B080/guz/simplifyGO_test/examples/random_BP/rds/clt_random_BP_1.rds")
mat = rl[[1]]
cl = binary_cut(mat)
ht_clusters(mat, cl, draw_word_cloud = FALSE)
```

Multiple cutoff can be tried by `select_cutoff()` function. The function applies binary cut clustering with every cutoff and calculates the difference score, the number of clusters and the block mean values for each clustering. Finally it produces a plot with the three metrics against different cutoff values.

In the following plot, basically a cutoff in the interval [0.72, 0.98] generates almost the same clusterings, and with cutoff lower than 0.71, the clusterings lead to more groups, which decreases the difference scores and increase the block mean values.

```
select_cutoff(mat, verbose = FALSE)
```

Next we show two clusterings with cutoff of 0.8 and 0.7 separately. When cutoff is set to 0.8, the clustering visually is the same as the heatmap we showed in the beginning of this document (where the cutoff is 0.85), and when cutoff is set to 0.7, some clusters are split into more sub-clusters.

For this case, we would say a cutoff of 0.8 is better than 0.7, as it generates higher difference score and the heatmap look cleaner.

```
cl = binary_cut(mat, cutoff = 0.8)
p1 = grid.grabExpr(ht_clusters(mat, cl, draw_word_cloud = FALSE, column_title = "cutoff = 0.8",
    show_heatmap_legend = FALSE))

cl = binary_cut(mat, cutoff = 0.7)
p2 = grid.grabExpr(ht_clusters(mat, cl, draw_word_cloud = FALSE, column_title = "cutoff = 0.7",
    show_heatmap_legend = FALSE))
grid.newpage()
pushViewport(viewport(x = 0.25, width = 0.5))
grid.draw(p1)
popViewport()
pushViewport(viewport(x = 0.75, width = 0.5))
grid.draw(p2)
popViewport()
```

There are some other scenarios where the cutoff needs to manually adjusted. The following example is based on one dataset from EBI Expression Atlas cohort, taking Disease Ontology as the gene sets. The heatmap below is based on the default cutoff of 0.85.

```
rl = readRDS("/icgc/dkfzlsdf/analysis/B080/guz/simplifyGO_test/examples/EBI_Expression_Atlas_DO/rds/clt_E-GEOD-22385_A-AFFY-37_g1_g2_DO.rds")
mat = rl[[1]]
cl = binary_cut(mat)
ht_clusters(mat, cl, draw_word_cloud = FALSE)
```

This is a very typical case where in the big clusters also exist multiple dense and small clusters. In this case, the cutoff can be fine-tuned according to what levels of similarities users want to keep.

If we run `select_cutoff()`, actually we can see the cutoff can lead to two different clusterings (cutoff in [0.83, 0.98] and in [0.67, 0.82]).

```
select_cutoff(mat, verbose = FALSE)
```

And if we set the cutof to some value lower than 0.83, the small dense clusters will be captured.

```
cl = binary_cut(mat, cutoff = 0.8)
ht_clusters(mat, cl, draw_word_cloud = FALSE, column_title = "cutoff = 0.8")
```
