## Supplementary file 10 for "*simplifyEnrichment*: an R/Bioconductor package for Clustering and Visualizing Functional Enrichment Results": suppl10_concordance.html (from another version of this preprint: version 1, 2020-10-28)

Supplementary file 10. Concordance score between two clusterings


### Supplementary file 10. Concordance score between two clusterings

###### Zuguang Gu

#### 2020-10-05

For the clusterings from two methods, first the cluster labels are adjusted. Denote \(\mathbf{s\_1}\) as the label vector for method 1, \(\mathbf{s\_2}\) as the label vector for method 2, and \(\mathbf{s\_1}\) as the reference labels, we apply `clue::solve_LSAP()` function to generate a mapping function \(m()\) between the two sets of labels to maximize \(\sum^n\_i I(s\_{1i}, m(s\_{2i}))\) where \(n\) is the length of \(\mathbf{s\_1}\) or \(\mathbf{s\_2}\). Denote the adjusted labels for the second method as \(s'\_{2i} = m(s\_{2i})\), the concordance between the two clusterings are calculated as:

\[ \frac{1}{n}\sum\_i^n I(s\_{1i}, s'\_{2i}) \]

where \(I(x, y)\) is an indicator function with value 1 if \(x = y\) and 0 if \(x \ne y\). A concordance of 1 means the two clusterings are identical.
